# A robust deep learning platform to predict CD8+ T-cell epitopes

**DOI:** 10.1101/2022.12.29.522182

**Authors:** Chloe H. Lee, Jaesung Huh, Paul R. Buckley, Myeongjun Jang, Mariana Pereira Pinho, Ricardo A. Fernandes, Agne Antanaviciute, Alison Simmons, Hashem Koohy

## Abstract

T-cells play a crucial role in the adaptive immune system by inducing an anti-tumour response, defending against pathogens, and maintaining tolerance against self-antigens, which has sparked interest in the development of T-cell-based vaccines and immunotherapies. Because screening antigens driving the T-cell response is currently low-throughput and laborious, computational methods for predicting CD8+ T-cell epitopes have emerged. However, most immunogenicity algorithms struggle to learn features of peptide immunogenicity from small datasets, suffer from HLA bias and are unable to reliably predict pathology-specific CD8+ T-cell epitopes. Therefore, we developed TRAP (T-cell recognition potential of HLA-I presented peptides), a robust deep learning platform for predicting CD8+ T-cell epitopes from MHC-I presented pathogenic and self-peptides. TRAP uses transfer learning, deep learning architecture and MHC binding information to make context-specific predictions of CD8+ T-cell epitopes. TRAP also detects low-confidence predictions for peptides that differ significantly from those in the training datasets to abstain from making incorrect predictions. To estimate the immunogenicity of pathogenic peptides with low-confidence predictions, we further developed a novel metric, RSAT (relative similarity to autoantigens and tumour-associated antigens), as a complementary to ‘dissimilarity to self’ from cancer studies. We used TRAP to identify epitopes from glioblastoma patients as well as SARS-CoV-2 peptides, and it outperformed other algorithms in both cancer and pathogenic settings. Thus, this study presents a novel computational platform for accurately predicting CD8+ T-cell epitopes to foster a better understanding of antigen-specific T-cell response and the development of effective clinical therapeutics.

**Highlights:**

- HLA bias and out-of-distribution problem are causes of poor performance of current state-of-the-art algorithms
- Transfer learning, deep learning architecture, context-specific and HLA-generalised approaches improve CD8+ T-cell epitope prediction
- TRAP reports degree of correctness to improve reliability of the prediction
- A novel metric termed RSAT estimates immunogenicity of pathogenic peptides, as a complementary to ‘dissimilarity to self’ from cancer studies

## Introduction

T-cells are essential for eliminating intracellular infections, triggering anti-tumour response as well as developing an immunological memory. Their ability to induce antigen-directed cytotoxicity has proven instrumental in fighting diseases, as evidenced by checkpoint blockade immunotherapy, adoptive cellular therapy and cancer vaccinology^1–3^. With the growing interest in identifying the cognate antigens of antigen-specific T-cells, many efforts have been made to improve experimental and computational methods for screening, predicting or characterising T-cell epitopes. However, current experimental approaches for identifying T-cell targets are labour-intensive, low-throughput and expensive^4–6^, and computational methods are still in their infancy^7^.

An effective antigen-specific CD8+ T-cell response to exogenous pathogens or endogenous threats relies on tightly regulated processing and presentation of antigenic peptides by class I MHCs, followed by recognition of the peptide-MHC (pMHC) by cognate CD8+ T-cells. Therefore, immunogenic peptides encompass features associated with MHC presentation *and* T-cell recognition^8^. Among these, it has been demonstrated that features attributed to MHC presentation are more prominent than those attributed to TCR recognition, with strongly conserved motifs at anchor positions being one such feature^9,10^. Indeed, recent cutting-edge models^11,12^, such as the widely used NetMHCpan^12^, have demonstrated impressive performance in predicting MHC presentation on certain alleles. On the other hand, the dual nature of the peptide-specific TCR recognition interface, comprised of both peptide and MHC, makes predicting the interaction between TCR and pMHC uniquely challenging. In addition, the scarcity of peptides tested by T-cell assays, as well as the lack of true negative datasets (i.e. presented but not recognized by T-cells), hampers our understanding of the properties underlying T-cell recognition.

Despite these challenges, a plethora of computational models have been developed to aid in the prediction of T-cell targets and to foster a better understanding of the characteristics underpinning peptide immunogenicity^8,13–17^. These models utilise features such as amino acids at contact position, large and aromatic side chains^13^, hydrophobicity^17–20^, peptide-MHC binding affinity and stability^16^ as correlates to T-cell recognition. Specifically for cancer neoepitopes, agretopicity (i.e. the ratio of binding affinity between neoepitope and wild-type counterpart), foreignness score (i.e. similarity of neoepitope to previously characterised epitopes)^8^ and dissimilarity to human proteome^21,22^ were found to indicate T-cell recognition.

However, we previously found that cancer and pathogenic epitopes often do not share the same immunogenicity features, which may differ in directionality or magnitude^23^. In addition to the context-specific differences, other factors, such as limited training data and a highly diverse T-cell receptor (TCR) repertoire contributed to the difficulty of predicting T-cell recognition potential. As a result, many existing models perform poorly against both cancer and emerging viral pathogens^7^, and progress in improving performance appears to be incremental, suggesting that there are still remaining challenges in predicting T-cell epitopes.

Over the last decade, deep learning and natural language processing (NLP) have transformed biomedical research and offered breakthrough discoveries^24^. Because of their ability to extract complex patterns from large amounts of data, deep neural network (DNN) models have been used for predicting peptide-HLA binding^12,25,26^ as well as TCR specificity^27–29^. Furthermore, transformer-based pre-trained language models (PLMs) have advanced the prediction of protein structure and function^30–33^ by combining the power of transformers, self-supervised learning and transfer learning^34–36^. Indeed, as a solution to data constraints, these protein PLMs, which contain knowledge learned from a large volume of protein sequences, could serve as an additional source of information for related downstream tasks.

In addition, more research has recently been conducted on providing reliable predictions for safety-critical applications. Most of DNN models are trained under the assumption that test data distribution will be similar to the training data distribution. However, when used in real-world tasks, out-of-distribution (OOD) examples that deviate from the training data are common^37,38^, resulting in a significant drop in model performance^39–41^. While this may be acceptable for some applications such as movie recommendations, it can be disastrous in safety-critical applications, such as medical diagnosis^42,43^. Therefore, the ability to identify OOD inputs and respond appropriately, whether by abstaining, requesting human intervention or gathering additional information, has become critical^42^. Recently, several methods for estimating the degree of correctness have been proposed, and have been successfully applied in the Natural language Inference (NLI) and/or OOD datasets^44–48^. One of the major challenges in immunogenicity prediction has been the limited data and OOD generalisation problem for peptides derived from different hosts, organisms and diseases. As such, the OOD detection module would facilitate making reliable predictions on a real-world set of peptides that are often highly diverse and heterogeneous.

Here, we present TRAP (<u>T</u>-cell <u>r</u>ecognition potential of HL<u>A</u>-I presented <u>p</u>eptides), a deep learning-based platform that addresses the current limitations and effectively captures T-cell recognition motifs from HLA-I presented pathogenic or self-peptides. Novel strategies were implemented, such as a) building separate models for pathogenic and self-peptides to account for divergent immunogenicity-related features, b) using transfer learning to deliver amino acid embeddings from pre-trained large-scale protein language models, c) capturing T-cell recognition motifs with a deep learning architecture, and d) detecting low confidence predictions to abstain from making incorrect predictions. We further developed RSAT (relative similarity to autoantigens and tumour-associated antigens) to estimate the immunogenicity of pathogenic peptides when they are abstained due to low-confidence predictions. The TRAP was then used to identify cancer neoepitopes from glioblastoma patients and showed superior performance to other methods. While many immunogenicity algorithms are based on MHC binding, TRAP goes one step further by predicting T-cell targets from MHC-I ligands. This novel workflow will enable more accurate identification of CD8+ T-cell epitopes, facilitating the development of effective vaccines and therapeutics.

## Results

### TRAP: A robust deep learning platform to predict CD8+ T-cell recognition of MHC-I presented pathogenic and self-peptides

We present TRAP as a comprehensive platform for predicting CD8+ T-cell immunogenicity of HLA-I presented pathogenic and self-peptides (Figure 1A).

**Figure 1.**
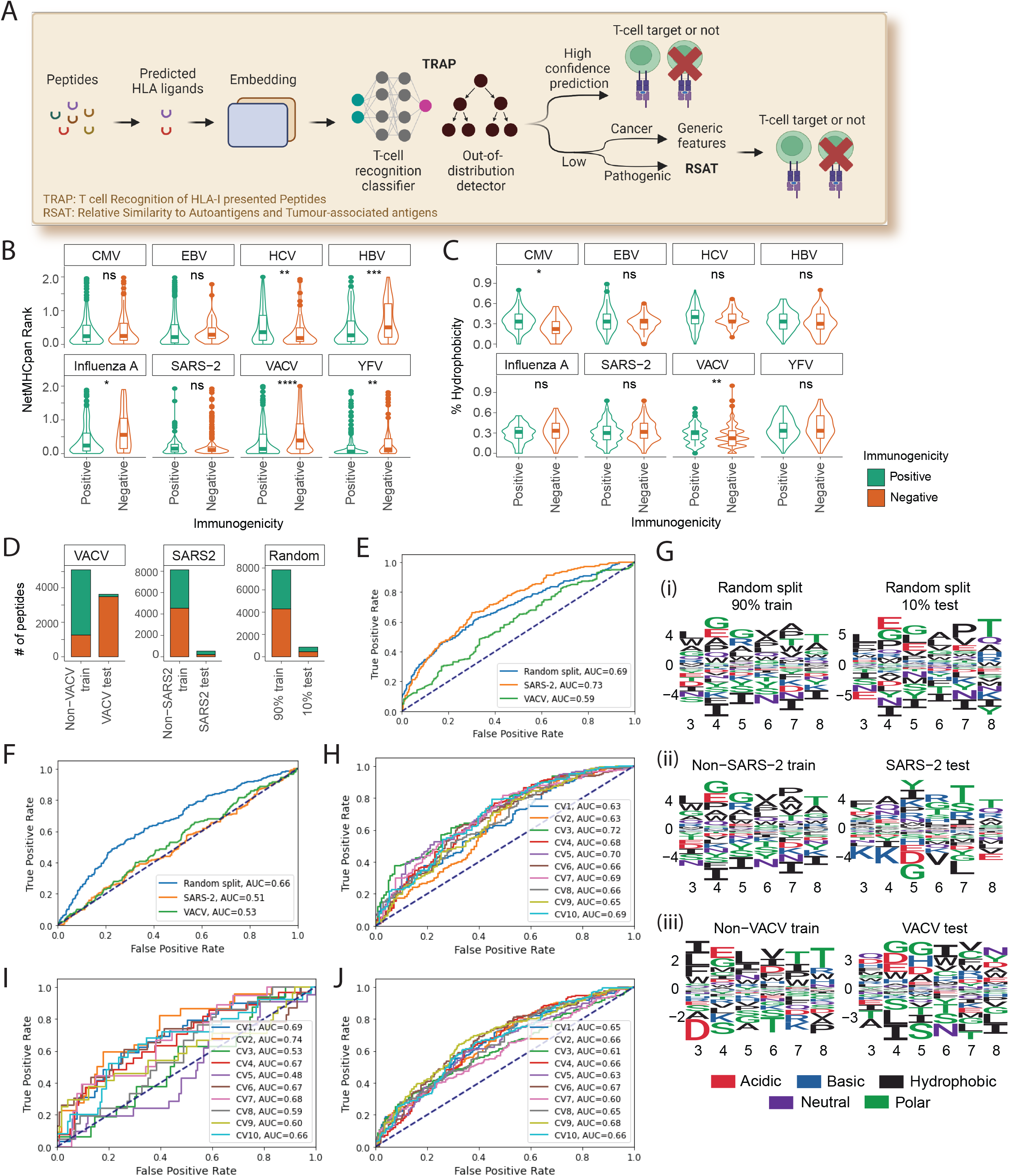
Schematic of TRAP workflow and cross-species variation in T cell recognition features. A. Schematic diagram of TRAP (T-cell recognition potential of HLA-I presented peptides), a robust deep learning-based workflow to predict CD8+ T-cell epitopes from MHC-I presented pathogenic or self-peptides. Once peptides have been predicted by NetMHCpan to bind HLA alleles, the TRAP uses the peptide sequence and NetMHCpan rank scores as inputs to predict the immunogenicity of the peptide with the respective HLA binding affinity. The TRAP workflow will output TRAP prediction score along with confidence in its prediction. If the prediction is detected to have a low confidence, we recommend predicting cancer neoepitopes using TESLA^52^, which is known to use more general features such as agretopicity and dissimilarity to self-proteome, and pathogenic peptides with RSAT (Relative Similarity to Autoantigens or Tumour-associated antigens). B-C. Distribution of MHC binding rank scores predicted by NetMHCpan (B) and hydrophobicity (C) for peptides derived from different pathogenic species. CMV: Cytomegalovirus; EBV: Epstein-Barr Virus; HCV: Hepatitis C Virus; HBV: Hepatitis B Virus; SARS-2: SARS-CoV-2; VACV: Vaccinia Virus; YFV: Yellow Fever Virus. D. Statistics of peptides in cross-species dataset (i.e. non-vaccinia virus (non-VACV) peptides for training and VACV peptides for testing, non-SARS-CoV-2 (non-SARS2) peptides for training and SARS-2 peptides for testing), and data randomly divided into 90% train and 10% test, as a resemblance of 10-fold cross-validation. E-F. Models trained using cross-species datasets could not effectively predict the immunogenicity of peptides derived from unseen pathogens. ROC-AUC curves of XGBOOST classifiers on training data by 10-fold cross-validations - on 90% data, non-SARS-2 and non-VACV peptides (E). ROC-AUC curves of XGBOOST classifiers on test datasets - 10% data, SARS2 and VACV peptides (F). G. Sequence logo of amino acids enriched in epitopes (Positive) compared to non-epitopes (Negative) in contact positions for randomly split data (i), SARS-2 data (ii) and vaccinia virus data (iii). H-I. High performance may be a reflection of HLA bias. ROC curve of DeepImmuno algorithm on single HLA allele, for peptides bound on HLA-A*02:01 (H) or HLA-A*24:02 (I). J. Performance of DeepImmuno algorithm on per-HLA down-sampled dataset i.e. the number of peptides has been down

In this workflow, peptide sequences of 9-10 amino acids in length are predicted to bind HLAs. The predicted HLA-I ligands are encoded using amino acid embeddings derived from protein transformer-based pretrained language models (PLMs). TRAP then employs a 1D convolutional neural network to extract T-cell recognition motifs, which are then combined with MHC binding rank score and hydrophobicity to predict peptide immunogenicity. Following that, TRAP includes a decision tree classifier to detect low-confidence predictions to improve accuracy. The immunogenicity of pathogenic peptides with low-confidence predictions can be predicted using a novel metric called Relative Similarity to Autoantigens or Tumour-associated antigens (RSAT), which is developed as a complement to “dissimilarity to self” from cancer studies.

TRAP has been developed as a user-friendly web application __, where users can input their peptide list and select the model of interest (pathogenic or self-antigen), and the application will compute the prediction scores along with its confidence (Supplementary Figure 1). In the following sections, we will explain the rationales behind the model architecture and strategies for overcoming the current constraints and improving model performance.

### Out-of-distribution uncertainty and HLA bias result in poor performance

We previously reported that the existing immunogenicity algorithms showed suboptimal performance in predicting epitopes from both cancer and an emerging viral pathogen^7^. We then attributed the poor performance to divergent discriminative features between cancer neoepitopes and pathogenic epitopes in directionality or magnitudes^7,23^.

Here, we investigated additional sources of suboptimal performance to aid in the development of accurate, robust and biologically meaningful classifiers. We retrieved peptides of lengths 9-10aa, characterised by T-cell assays from IEDB and retained predicted MHC binder from NetMHCpan 4.0, resulting in 5,093 immunogenic (‘Positive’) and 6,628 non-immunogenic (‘Negative’) peptides (named PeptideTcell data, Methods) (Supplementary Figure 2A). Notably, no peptides may be considered truly non-immunogenic as they may be recognized by at least one TCR in the repertoire under the right physiological conditions. However, there is a continuum of immunogenicity potential where some peptides are likely to trigger greater T-cell response and/or are able to bind numerous TCRs. Therefore, we designated the ‘Negatives’ as peptides with very low putative immunogenicity potential, defined as peptides characterised negative in more than three T-cell assays.

First, we evaluated the extent to which known hallmarks of immunogenicity, such as MHC binding affinity^12,16^ and percentage of hydrophobic amino acids in peptide sequence^20^, can be used to identify immunogenic peptides from different species. Notably, these features have been used as predictors in models such as TESLA^8^, iPred^17^, NetTepi^16^ and GAO^49^. Although two features could effectively distinguish epitopes from non-epitopes at a macroscopic level, where epitopes had higher MHC binding rank (-log transformed) and hydrophobicity with medium effect sizes (Supplementary Figure 2B-C), they no longer showed statistical significance at the species-level (Figure 1B-C), which was caused in part by the small sample size. For example, hydrophobicity alone could not discriminate SARS-CoV-2 epitopes from non-epitopes.

Second, we assessed the extent to which cross-species peptides (i.e. peptides derived from other species) can predict epitopes of unseen pathogens. We divided PeptideTcell data randomly or in a cross-species manner: i) random split to 90% train vs. 10% test (i.e. representation of 10-fold cross-validation), or ii) cross-species split to Non-SARS-CoV-2 (Non-SARS-2) train vs. SARS-2 test, and Non-vaccinia virus (Non-VACV) train vs. VACV test (Figure 1D). We then compared the performance of XGBOOST classifiers on these datasets (Method). While no substantial difference was observed during training (Figure 1E), the cross-species model showed lower accuracy and higher root mean squared error (RMSE) on test datasets (Figure 1F). This implied that cross-species peptides may not share common features in predicting immunogenicity (i.e. predictive features in training and test data are likely to be different which is known as an out-of-distribution generalisation problem), resulting in limited accuracy on unseen pathogens.

To support this, we generated differential position specific scoring matrices (dPSSMs) to compare immunogenicity-related sequence patterns between train and test datasets (Methods). While random split data shared similar patterns between train and test datasets, such as enrichment of L, G/E, G, hydrophobic resides and T on P3-P8 (Figure 1Gi), cross-species data showed low homology (Figure 1Gii-iii), and dPSSM scores failed to predict immunogenicity on cross-species test datasets (Supplementary Figure 2D, Method).

Third, we observed that high reported performance from the latest models incorporating peptide-HLA pairs might be driven by HLA bias from an imbalanced dataset. Recently, deep learning models incorporating peptide-HLA pairs reported ROC-AUC of ~0.85 by 10-fold cross-validation^15^ (Supplementary Figure 2E-F). While they reported the highest accuracy to date, we observed poor performance on SARS-CoV-2 peptides in our benchmarking study^7^. Here, we hypothesized that this poor agreement may be due to differences in datasets and investigated the cause of disagreement.

First, we evaluated the ability of the model to discriminate Positive vs. Negative bound on the same HLA allele. We trained and tested models in single HLA-level on HLA-A*02:01 and HLA-A*24:02, which had the highest numbers of characterised peptides per HLA. We then compared them to the model trained using the same number of randomly sampled pMHCs (Supplementary Figure 2G). Here, the model showed marginally better than random performance on the balanced HLA-A*02:01 dataset (Figure 1H), and high variation on the relatively imbalanced and smaller HLA-A*24:02 dataset (Figure 1I). We further found that HLA-balanced data substantially reduces classifier performance (Supplementary Figure 2H, Figure 1J), suggesting that current models are skewed towards classifying for certain over- or under-represented HLA alleles and their reported performance does not reflect their real-world accuracy.

### Mitigate HLA bias by employing peptide sequences at TCR contact positions

Previous studies reported the contribution of anchor positions (i.e. position 2 (P2) and P9 of 9aa peptide) for MHC binding^50^ and contact positions (i.e. P3-P8) in T-cell recognition^13^. Correspondingly, HLA supertypes drove the clustering of peptides at anchor positions (Figure 2A) and TCR specificities at contact positions (Figure 2B-C, Supplementary Figure 3A-B) on our data, with peptides bound by the same TCR showing conserved motifs (Figure 2D). However, because of the strong conserved pattern in anchor positions, the HLA supertype not only drove the clustering of peptides at anchor positions but also acted as the strongest covariate driving the clustering of peptides in full sequence (Figure 2E, Supplementary Figure 3C). With such a strong conserved pattern, MHC binding features may dominate immunogenicity predictions when the full peptide sequence is used for model training. In fact, we observed that some of the existing immunogenicity classifiers were more capable of predicting dominant HLA type (i.e. whether peptides were bound to HLA-A*02:01 or not) than peptide immunogenicity^7^. As we aimed to predict T-cell recognition potential once peptides are bound to HLA alleles, we incorporated contact positions only i.e. positions 3-8 (P3-P8) of 9aa peptide and P3-P9 of 10aa peptides, not only to focus on T-cell recognition patterns, but also to rule out the need to balance the number of epitopes vs. non-epitopes by the HLA supertypes for model development.

**Figure 2.**
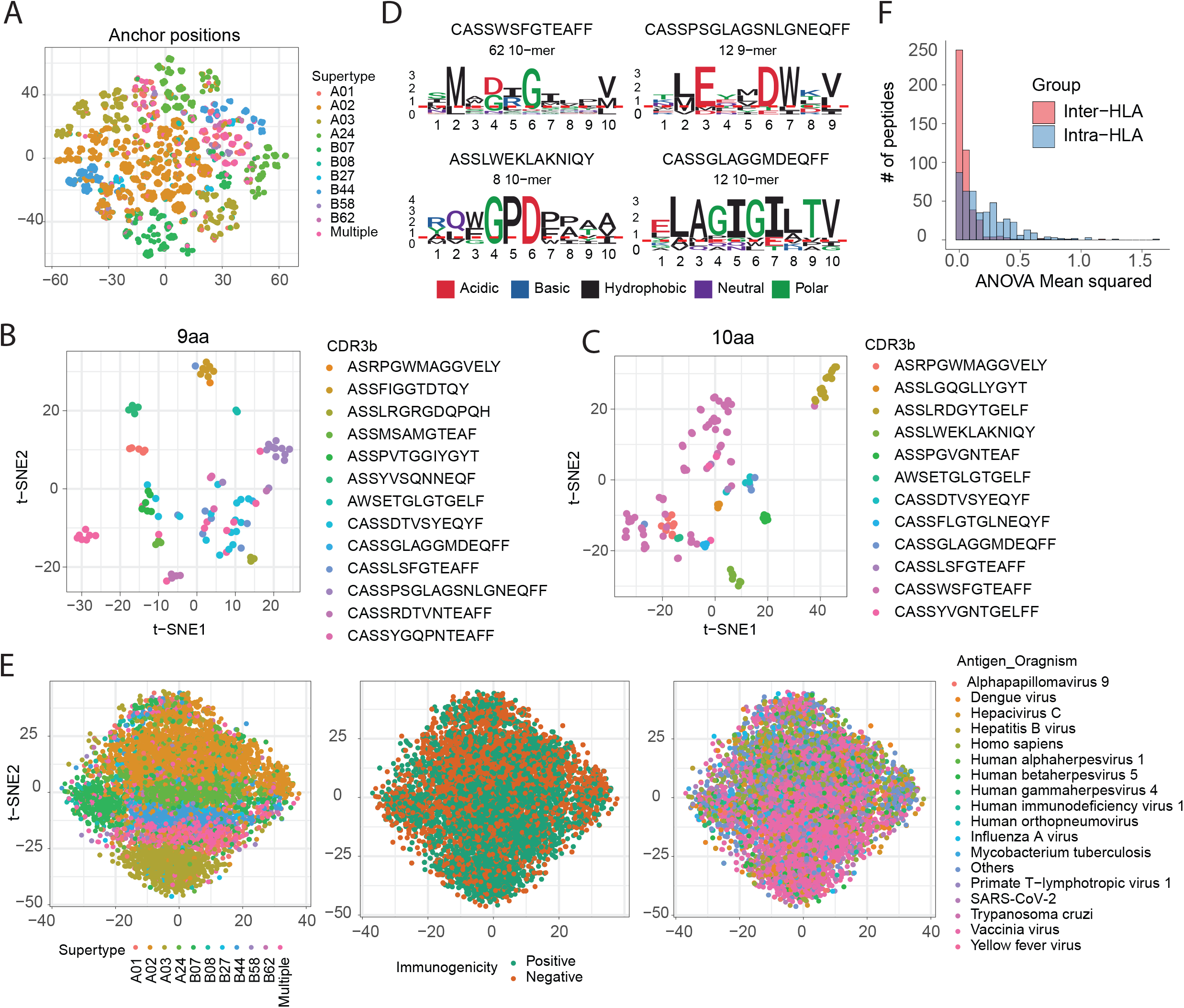
Effect of anchor and contact positions on peptide immunogenicity. A. t-SNE embedding of peptides, coloured by HLA supertypes. Peptides at anchor positions (i.e. P2 and P9 of 9aa peptides) were represented by amino acid descriptors (aaDescriptors) and all amino acids across the peptide sequence were averaged to compute peptide-wide aaDescriptors. Each peptide was represented by positional (only at anchor positions) and peptide-wide aaDescriptors. B-C. t-SNE embedding of peptide-wide descriptors and amino acids at contact positions, coloured by cognate TCRs for 9aa peptides (B) and 10aa peptides (C). D. Sequence logos of peptides recognised by the same TCR, coloured by the physicochemical properties. Notably, these are examples of TCRs having the most cognate peptides in the database. However, except for the first TCR, the low number of peptides may limit statistical confidence in representing sequence conservation. E. t-SNE embeddings of peptide-wide descriptors and amino acids at all positions, coloured by HLA supertype (left), immunogenicity (middle) and species of origin (right). F. Distribution of intra-HLA and inter-HLA variation for peptides having different peptide-HLA entries. The variation described by ANOVA means squared.

To take such an approach, we first investigated whether neglecting HLA information would not result in a significant loss of information when predicting T-cell recognition potential of MHC-presented peptides from the current dataset. First, a novel metric was developed for converting qualitative measurements (i.e. Positive or Negative) to a quantitative ‘positivity score’ that reflects confidence in the immunogenicity (Methods). Briefly, the metric took into account a) the number of experiments conducted (Supplementary Figure 4A), b) the percentage tested positive (Supplementary Figure 4B) and c) the number of cognate TCRs where available (Supplementary Figure 4C) to compute the positivity score (Supplementary Figure 4D).

Using the positivity scores of each peptide-HLA entry, the extent of inter-HLA variation (i.e. variation across different HLA alleles) was compared to intra-HLA variation (i.e. variation within the same HLA allele) for peptides bound on multiple HLA alleles. The intra-HLA variation accounted for the majority of data variability, while inter-HLA variation contributed less (Figure 2F). This is because other biological (e.g. different effector cell, antigen presenting cells etc.) or technical (e.g. assay type, culture conditions, research groups etc.) factors were causing greater discrepancies in positivity score than HLA alleles (Supplementary Figure 4E-F). This implied that when it came to each peptide, there wasn’t much difference in the immunogenicity based on the HLAs to which it was bound, but when it came to HLAs, each HLA had a different pool of peptides, resulting in a different distribution of positive and negative peptides, resulting in HLA-bias. As a result, an HLA-generalised approach was adopted in which peptide-HLA binding information was used instead of the HLA allele itself.

### Deep learning architecture captures T-cell recognition motifs

As the potential causes of poor performance, previous sections discussed limited training dataset, HLA bias and out-of-distribution (OOD) generalisation problem. These issues were addressed in TRAP by a) using peptide sequence at contact positions, b) building separate models for pathogenic and self-peptides, c) encoding peptide sequences using amino acid properties derived from protein transformer-based pretrained language models (PLMs), d) devising one-dimensional convolutional neural network (1D CNN) architecture designed to capture T-cell recognition motifs and e) employing OOD detection module. These novel strategies enabled TRAP to offer more accurate and reliable predictions of CD8+ T-cell targets against cancer and viral diseases.

First, pathogenic and self-antigen datasets were prepared to build context-specific models (Methods). Briefly, the pathogenic dataset is a subset of the PeptideTcell data, comprising only pathogen-derived peptides (Supplementary Figure 5A). For the self-antigen dataset, autoantigens, tumour-associated antigens and cancer neoepitopes were retrieved from Cancer peptide database, dbPepNeo, IEDB, McPAS-TCR and NEPdb databases as epitopes, and benign HLA-I ligands expressed in thymus as non-epitopes (Supplementary Figure 5B-C). The HLA-I ligands (exclusively) expressed in the thymus were assumed to be involved in negative selection of T-cells, and thus no repertoire to recognise these peptides was expected. To the best of our knowledge, this is the first study to use the concept of negative thymic selection in classifying self-epitopes from non-epitopes.

Second, because peptides vary in lengths, we investigated the optimal padding strategy for aligning 6-mer with 7-mer contact position peptides. The predictive accuracies of simple dense layer classification, 1D CNN, 2D CNN and bidirectional recurrent neural network (biRNN) models were evaluated using pre- and post-padding strategies. No significant difference was observed across pathogenic and self-antigen data, with pre-padding achieving slightly better performance in self-antigen data on the BiRNN model (Figure 3A). Furthermore, the relative location of k-gram motifs was compared on 6-mer and 7-mer peptides, and many 3-gram motifs found at P3 of the 9aa peptide were found at P4 of the 10aa peptide (Figure 3B). Therefore, to align T-cell recognition hotspots with biological context, peptides of shorter length were pre-padded to align with longer peptides.

**Figure 3.**
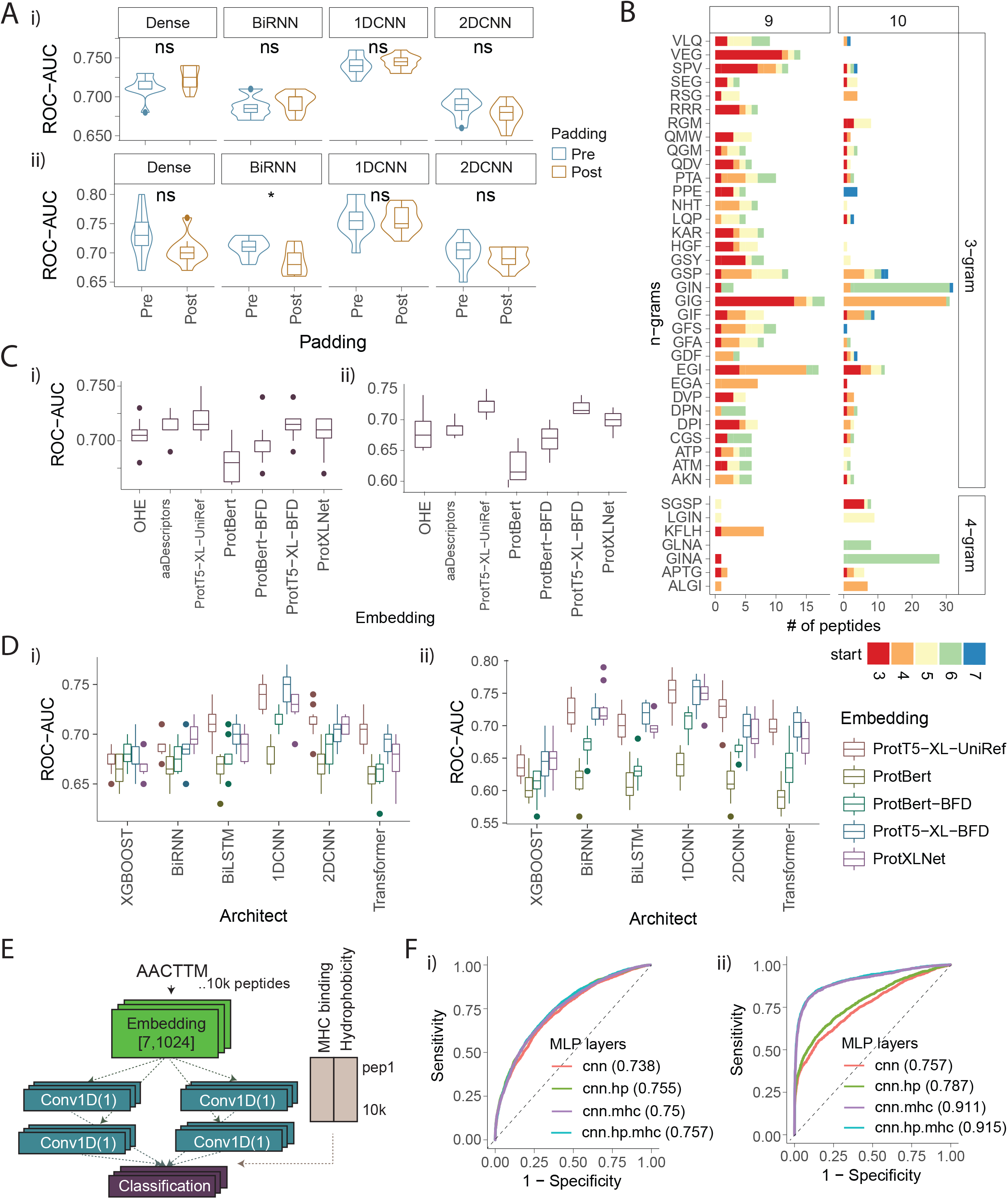
Optimise TRAP architecture. A. Performance of pre- and post-padding strategies. ROC-AUC values of respective deep learning models with pre- vs. post-padding strategies for pathogenic (i) and self-antigens (ii). Each point represents the ROC-AUC value from one round of 10-fold cross-validations. B. Example of n-grams found in both 9aa and 10aa peptides, coloured by the start position of the respective n-gram in the peptides. C. Comparing amino acid embedding strategies. ROC-AUC values of a single dense classification prediction using different encoding strategies, including one-hot encoding (OHE), amino acid descriptors (aaDescriptors) and embeddings from protein transformer-based PLMs on pathogenic peptides (i) and self-antigens (ii). Each point represents the ROC-AUC value from one round of 10-fold cross-validations. D. Comparing the performance of different deep learning architectures. ROC-AUC values comparing the performance of different machine learning and deep learning architectures using embeddings from 5 protein transformer-based PLMs for pathogenic (i) and self-peptides (ii). XGBOOST: an extreme gradient boosting. BiRNN: bidirectional recurrent neural network. BiLSTM: bidirectional Long short-term memory. 1DCNN: 1-dimensional convolutional neural network. 2DCNN: 2dimensional CNN. Each point represents ROC-AUC value from one round of 10-fold cross-validations. E. Schematic diagram of the model incorporating MHC binding and hydrophobicity. F. Adding hallmarks of the immunogenicity. ROC curve comparing the performance of models after adding MHC binding rank score and/or hydrophobicity to peptide sequence-based 1D CNN model for pathogenic (i) and self-peptides (ii).

Third, to address data limitation, we adopted transfer learning to encode peptides using amino acid properties derived from protein transformer-based pretrained language models (PLMs). The protein transformer-based PLMs were trained using millions to billions of protein sequences and carry the most representative 1024 embeddings, which describe the physicochemical, structural or electrostatic properties of amino acids in protein space^33^ (Supplementary Table 2). The performance of classification models were compared when peptides were encoded by one-hot-encoding (OHE), amino acid descriptors and embeddings from five protein transformer-based PLMs (Methods). ProtT5-XL-UniRef showed the highest performance, with an average ROC-AUC of 0.72 for pathogenic (Figure 3Ci) and 0.724 for self-antigen datasets (Figure 3Cii).

Fourth, the capability of different machine learning and deep learning architectures to classify immunogenicity potential was assessed. The performances of XGBOOST, BiRNN, bidirectional long short-term memory (BiLSTM), 1D CNN, 2D CNN and transformer models were compared. Briefly, RNNs or LSTMs are popular sequential models efficient in addressing sequential text data^51^, whereas CNNs are suitable for scanning across text or images and detecting local patterns by using ‘kernels’. 1D CNNs have widely been used for text and 2Ds for image classifications^52^. The 1D CNN model using ProtT5-XL-UniRef embeddings achieved the highest average ROC-AUC of 0.74 for pathogenic (Figure 3Di) and 0.75 for self-antigens (Figure 3Dii). We surmised that T-cell recognition motifs captured by the 1D CNN model have more predictive power than sequential relationship captured by the RNN-based models.

In addition to peptide sequences, we examined whether incorporating other hallmarks of immunogenicity, such as predicted MHC binding rank score^12,16^ and hydrophobicity^20^, can further improve the performance. Particularly, given that the current model only takes contact positions into account, adding MHC binding rank would allow the model to accommodate peptide-MHC binding information. This strategy has several advantages. First, it feeds the binding information with a minimal HLA-associated bias, circumventing the need to balance the training data by HLAs. Second, the most accurate binding information from NetMHCpan prediction can be incorporated without having to re-train the model using a massive peptide-HLA binding data. By integrating MHC binding rank and hydrophobicity as fully connected layers (i.e. Multilayer perceptrons, MLPs) and optimising hyperparameters (Figure 3E, Methods), performance was improved to an average ROC-AUC of 0.76 for pathogenic (Figure 3Fi) and 0.90 for self-antigens (Figure 3Fii). Of note, a large improvement in the self-antigen model was due to a greater difference in MHC binding rank between self-epitopes vs. non-epitopes, as a consequence of the relaxed MHC binding rank threshold (≤ 10 MHC binding rank) to account for as many self-epitopes.

Therefore, representing peptide sequences at contact positions by ProtT5-XL-UniRef amino acid embeddings, extracting T-cell recognition motifs by 1D CNN kernels, and adding MHC binding rank and hydrophobicity as MLPs could effectively achieve superior performance in classifying immunogenicity.

### Sequence patterns discriminating immunogenicity

Among different architectures, the 1D CNN model that captures local motifs achieved the best performance. Moreover, extensive screening of pMHC library against a single TCR revealed dominant hotspots or motifs in the cognate peptides^53,54^. We therefore set out to expand this observation and explore the enrichment of n-grams (i.e. contiguous sequence of n-amino acids)^55^ and position-specific k-mer motifs (i.e. contiguous or non-contiguous sequence of k amino acids at specific positions) in contact residues of pathogenic and self-peptides.

We first computed the ratio (in normalised frequency) of n-gram or positions-specific k-mer motifs between epitopes and non-epitopes (Methods). The top-ranking n-grams were GIG, GINA, GIF, LGIN, VEG and SGSP for pathogenic epitopes and SC, GIGI, IC, QC and CA for self-epitopes (Supplementary Figure 6C). In addition, top position-specific k-mer motifs were .EG.L., .E.IL. and .GIG… for pathogenic epitopes and GIG.., ..M.P., .G.GI.., and AGI…. for self-epitopes (Supplementary Table 3). Notably, GIG, which was previously associated with DMF5 TCR, was one of the top n-grams and position-specific k-mer motifs.

To identify sequence patterns enriched in both pathogenic and self-epitopes, we compared n-grams and position-specific k-mer motifs from each analysis. The 48 n-grams and total 298 position-specific k-mer motifs from 9 and 10aa peptides (Figure 4A) were shared between pathogenic and self-epitopes. These include GIG, GINA, GMP, ALGI and APTG n-grams (Figure 4B, Supplementary Figure 6D), and ….VP, W..P.., .G.GI.., and .GIG… positionspecific k-mer motifs (Supplementary Figure 6E-G).

**Figure 4.**
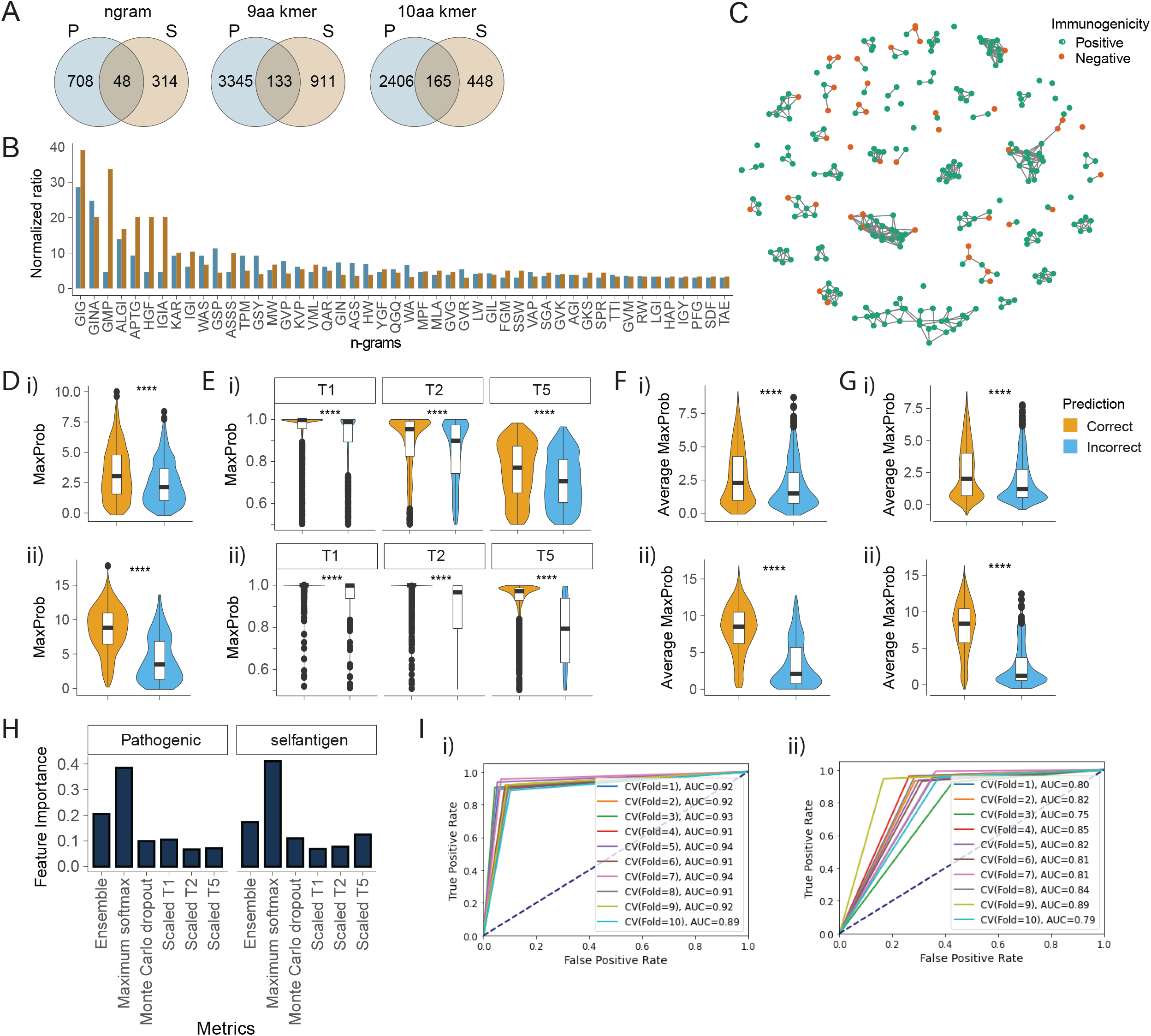
Out-of-distribution detection using calibration methods. A. Identifying motifs that are enriched in both pathogenic and self-epitopes. Venn diagram comparing the enriched n-grams (left) or position-specific k-mer motifs from 9aa (middle) or 10aa peptides (right) between pathogenic and self-peptides. B. Normalized enrichment ratio of shared n-grams enriched in both pathogenic and self-epitopes. C. Clusters of peptides with high sequence similarity in contact positions, demonstrated by pairwise global alignment scores. Network graph illustrating the pairwise global alignment score between 9aa pathogenic peptides. Shown are peptides having ≥22 alignment scores with ≥3 other peptides. D-G. Distribution of different calibration-based metrics on pathogenic (i) and self-(ii) peptides, discriminating correctly vs. incorrectly predicted peptides. The calibration-based metrics include maximum softmax probability (MaxProb) (D), temperature scaling using different levels of temperatures that scale the logit values (E), the average maximum softmax probability (average MaxProb) of 10 ensembled models (F) and the average MaxProb of models containing Monte Carlo dropouts (G). The Monte Carlo models were reiterated 100 times with stochastic dropouts of 0.6. T: temperature. Statistical significance by p-values from Student’s t-test. ns: non-significant. H. Feature importance of different calibration metrics on decision tree classifier that aims to discriminate correctly predicted peptides. Ensemble: average MaxProb of 10 ensembled models. I. ROC curve illustrating the performance of the out-of-distribution (OOD) decision tree classifiers using MaxProb and average MaxProb from ensembled models on pathogenic (i) and self-peptides (ii).

Furthermore, we computed pairwise global-alignment scores across pathogenic peptides to identify clusters of epitopes sharing high sequence homology (Method). Plotting highly similar peptides revealed clusters dominated by Positive peptides (Figure 4C). These clusters contained peptides having patterns, such as YNTI… FG(Y/F)PV(F/Y). F(E/D)(K/R)S.., G.GW.. .DR.WC, GPG.R.P, P.DFFP., .GIGI.., .DRGM., VALG.NA. and .LGLNA. (Supplementary Figure 5H-I).

Our analyses therefore suggest that TCR contact positions of immunogenic peptides exhibit preferences towards presenting certain motifs while disfavouring others. We also observed similarities and differences between pathogenic and self-peptides. The toolkits for analysing differential sequence patterns between peptides of the same length are developed as a R diffSeqPatterns package. We believe these shortlisted sequence patterns can be target for functional validations in identifying immune hotspots.

### Detect low-confidence predictions to improve robustness

While the DNN models could achieve higher accuracies than existing models, they are nonetheless far from making a perfect prediction. Partially due to the limited number of peptides characterised to fill the full combinatorial peptide space, the performance of sequence-based model drops substantially in predicting peptides non-homologous to those of training data, called the out-of-distribution (OOD) generalisation problem^43^. In contexts where incorrect predictions can have severe consequences such as in healthcare or security, using model uncertainty to decide when to trust or abstain the prediction can facilitate rejecting false predictions and improving model accuracy.

There are generally two types of OODs: background shift (i.e. a shift in population-level features that do not depend on classification labels) and semantic shift (i.e. a shift in features that are correlated with the context and label)^45^. In previous NLP studies, density or proximity-based models were found to be better at detecting background shifts, while calibration methods (i.e. using model’s prediction confidence) performed better at semantic shifts. In our study, the greatest OOD came from peptides originating from different species that have moderately different features associated with immunogenicity (i.e. semantic features). This aligns with previous findings that the majority of OOD stems from the semantic shift and thus calibration methods generally outperform proximity-based methods (i.e. autoencoder reconstruction)^45^.

Here, several OOD detection methods were tested to detect low-confidence predictions. Different autoencoder architectures, which are often used for anomaly detection, were first investigated^56^ (Supplementary Figure 7A-F, Methods). However, we observed no significant difference in reconstruction loss between correctly and incorrectly predicted peptides, implying that the autoencoder-based methods cannot effectively identify OOD inputs.

Therefore, calibration methods such as maximum softmax probability (MaxProb), temperature scaling, average MaxProb from an ensemble of 10 models (called Ensemble) and Monte Carlo dropout were tested, because they were reported to effectively detect semantic shifts (Methods). We observed that all four methods could significantly differentiate correct vs. incorrect predictions for both pathogenic and self-antigens (Figure 4D-G). To evaluate the predictive power of these metrics, we trained a decision tree classifier that takes the score of the calibration methods as an input and predicts whether the input is an OOD sample. We observed that MaxProb and Ensemble had the highest feature importance, i.e. possess the best predictive power (Figure 4H). They could achieve comparable performance to the model incorporating all four metrics with average ROC-AUC of 0.923 for pathogenic and 0.806 for self-antigens (Figure 4I). With these, a MaxProb and Ensemble-based OOD detection module was introduced downstream of 1D CNN model prediction to report peptides that are likely to have a correct prediction.

### Relative similarity to autoantigens or tumour-associated antigens (RSAT) as a novel feature of pathogenic peptide immunogenicity

We and others^23,57^ have shown that some of the highly predictive metrics found from cancer neoepitope studies, such as ‘dissimilarity to self’, may not be applicable to pathogenic peptides. To address this, we present an alternative solution, termed a relative similarity to autoantigens or tumour-associated antigens (RSAT), to estimate the immunogenicity potential of pathogenic peptides. For pathogenic peptides that suffer from low confidence prediction, RSAT can provide an additional estimate of immunogenicity potential.

The ‘dissimilarity to self’ stems from the paradigm of negative selection where T-cells that bind strongly to self-peptides should have been negatively selected and thus no T-cell repertoire should be present to bind peptides homologous to self-proteome^21,22^ (Figure 5A). However, there is another side of the story where T-cells that have low or moderate binding to self-peptides should have been positively selected. In fact, recent studies attributed the inability of the immune system to recognise a large number of pathogenic peptides – most of which are highly dissimilar to human proteome – to the mechanism of positive selection where only T-cells bound by low or moderately binding self-peptides survive to trigger an immune response^57^. Therefore, we hypothesised that pathogenic peptides homologous to immunogenic self-peptides, such as autoantigens or tumour-associated antigens, may be more likely to trigger an immune response, and assessed whether relative similarity to autoantigens or tumour-associated antigens compared to reference human proteome can be a predictor of immunogenicity for pathogenic peptides.

**Figure 5.**
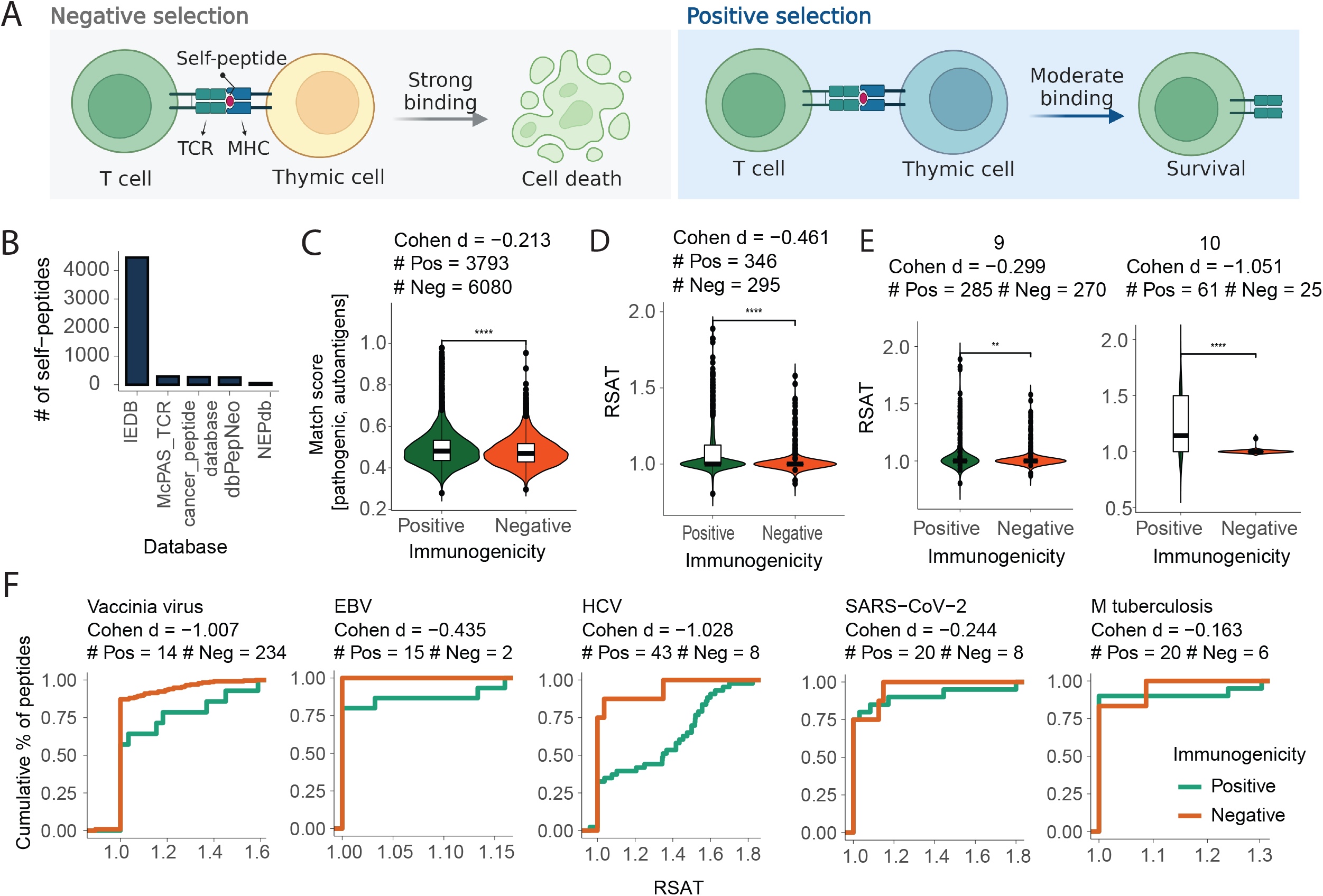
Relative similarity to autoantigens and tumour-associated antigens (RSAT). A. Schematic of positive and negative selection of T-cells in thymus. B. Number of self-peptides, such as cancer neoepitopes, autoantigens and tumour-associated antigens, retrieved from different databases. C. Distribution of Match score between pathogenic peptides and best self-peptide counterparts. Contains the Cohen’s d value showing the effect sizes differentiating positives versus negatives as well as the number of positive and negative peptides in each species. Cohen’s d values describe effective sizes, which are small (d = 0.2), medium (d = 0.5) and large (0.8). D. RSAT of pathogenic peptides that have comparable self-antigen counterparts (by match score ≥ 0.6). E-F. RSAT can effectively discriminate pathogenic epitopes from non-epitopes of varying lengths and species. E. Distribution of RSAT for pathogenic peptides having 9aa (left) and 10aa (right) in length. F. RSAT of peptides derived from different pathogens.

To test our hypothesis, 5023 unique Autoimmunity, Allergy or Tumour-associated antigens (AATs) were retrieved from NEPdb^58^, dbPepNeo^59^, McPAS-TCR^60^, IEDB^61^ and cancer antigenic peptide database (https://caped.icp.ucl.ac.be) (Figure 5B). We then computed the similarity between pathogenic peptides and AATs, and only retained pathogenic peptides having comparable AAT counterparts (Methods). On these pathogenic peptides, we then computed the similarity between pathogenic peptide and AAT’s closest human proteome counterparts, and the ratio between the two i.e. relative similarity of the pathogenic peptide to <u>a</u>utoantigens or <u>t</u>umour-associated antigens (named RSAT).

From pathogenic data, 346/3793 pathogenic epitopes and 295/6080 non-epitopes had comparable AAT counterparts by a threshold match score of 0.6 (Method, Figure 5C). The RSAT was computed on these peptides and found a significant difference between epitopes vs. non-epitopes (Figure 5D) across peptide lengths 9 and 10aa (Figure 5E), and across different pathogenic species (Figure 5F), indicating that RSAT could effectively classify peptides of varying lengths and pathogen species. Despite the fact that each species had a limited number of peptides, epitopes had higher RSAT values with low (SARS-CoV-2 and M.tuberculosis), medium (EBV) or high (HCV and vaccinia virus) effect sizes.

Here, we developed a novel metric, RSAT, to estimate the immune potential of pathogenic peptides based on relative similarity to known auto- or tumour-associated epitopes. We demonstrated that RSAT can effectively discriminate epitopes of different lengths and pathogens. The RSAT is available as a separate module in the TRAP GitHub repository and is recommended to be used in conjunction with TRAP when the pathogenic peptides have low prediction confidence. We appreciate that the low number of Autoimmunity, Allergy or Tumour-associated antigens (AATs) limits the use of RSAT on a broader range of peptides. However, we envision that as data becomes more abundant, RSAT can become more popular.

### Benchmark TRAP performance to state-of-the-art algorithms

The performance of TRAP was benchmarked against existing immunogenicity models, namely NetTepi, IEDB, PRIME, DeepImmuno and TESLA by 10-fold cross-validation on the same datasets (Method). Due to the nature of the existing models, HLA-agnostic predictions were performed for IEDB, iPred, NetTepi PRIME and Repitope, and HLA-restrictive prediction for DeepImmuno. TRAP was able to make both HLA-agnostic and restrictive predictions by incorporating the peptide sequence at contact positions and MHC as a rank score. The HLA-restrictive prediction was made using HLA-balanced data where the number of epitopes and non-epitopes were balanced per HLA to validate their prediction irrespective of HLA-associated bias.

TRAP was the best self-antigen model, achieving ROC-AUC of 0.931 for HLA-agnostic and 0.703 for HLA-restrictive predictions (Figure 6A). It was also one of the best pathogenic models, with ROC-AUC of 0.751 for HLA-agnostic and 0.709 for HLA-restrictive predictions (Figure 6B).

**Figure 6.**
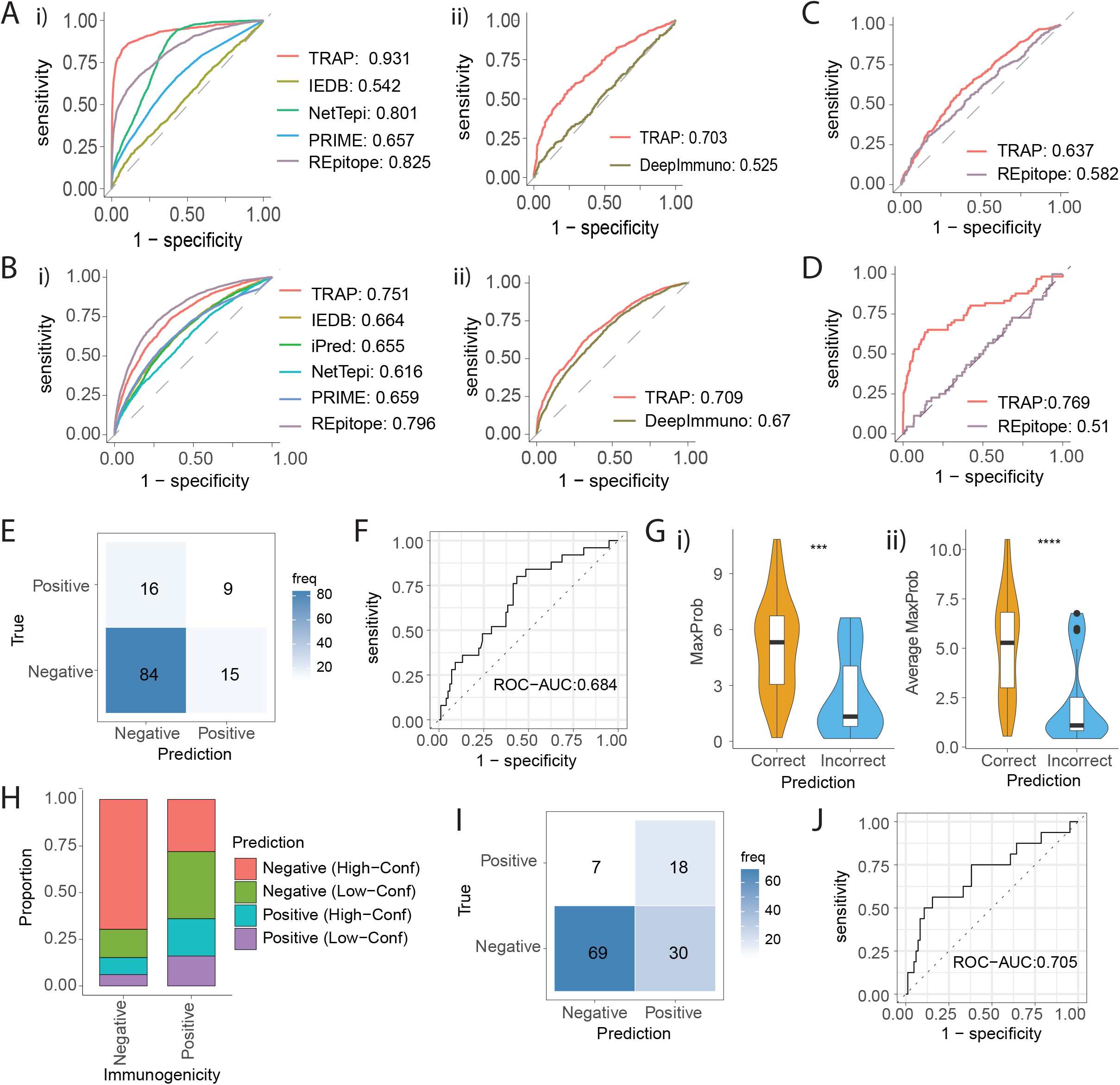
Benchmark TRAP performance and use it to identify tumour antigens in glioblastoma. A-B. ROC curve comparing the performance of TRAP with existing models, such as IEDB, iPred, NetTepi, PRIME, Repitope and DeepImmuno for self- (A) and pathogenic peptides (B). Due to the nature of these models, IEDB, iPred, NetTepi PRIME and Repitope were predicted in an HLA-agnostic manner (i.e. by peptide sequence only) (i), and DeepImmuno predicted in an HLA-restrictive manner (i.e. by peptide-HLA) using HLA-balanced data (ii). C. ROC curve comparing the performance of TRAP and Repitope that have been trained using 1511 Non-Wuhan SARS-CoV-2 peptides by 10-fold cross-validations. D. ROC-curve comparing the performance of Non-Wuhan SARS-CoV-2 trained TRAP and Repitope models in predicting 66 Wuhan SARS-CoV-2 peptides. E-J Application of TRAP in identifying glioblastoma neoepitopes. E. Confusion matrix of GBM cancer neoepitope prediction using self-antigen TRAP model in reference to T cell assay readout. F. ROC curve of TRAP performance on GBM dataset. G. MaxProb (i) and average MaxProb of 10 ensembled models (ii) for GBM peptides predicted to be non-immunogenic. H. The proportion of GBM epitopes (Positive) and non-epitopes (Negatives) predicted Positive or Negative with high or low confidence. The confidence has been predicted by the self-antigen out-of-distribution decision tree classifier. I. ROC curve of TRAP performance taking confidence into account, in which peptides that were predicted Negative with low confidence were removed to be considered as the potential epitope candidates. J. Confusion matrix of TRAP prediction after taking the confidence into account, in which peptides that were predicted Negative with low confidence were included as Positive.

While Repitope slightly outperformed TRAP in pathogenic datasets, the robustness of its predictions was assessed on emerging pathogens. The coronavirus TRAP and Repitope models were trained using 1511 Non-Wuhan SARS-CoV-2 peptides by 10-fold cross-validations (Figure 6C), which were then used to predict the immunogenicity of 66 Wuhan SARS-CoV-2 peptides (Figure 6D). Here, TRAP was better at extracting immunogenicity-related features from limited data of emerging pathogens, and translating them onto related species (Figure 6D).

Therefore, TRAP can predict CD8+ T-cell epitopes in both HLA-agnostic and restrictive contexts, and performed better in both. The TRAP can also make accurate predictions about emerging pathogens and is suitable for shortlisting therapeutic candidates when data is limited.

### Application of TRAP to identify glioblastoma neoantigens

Here, we demonstrate how the TRAP can be used for identifying cancer neoepitopes from glioblastoma patients. We previously sequenced the tumours of four HLA-A2 glioblastoma patients and shortlisted the cancer neoepitopes by using an in-house version of MuPeXI^62^ codenamed TUNAPASTA v0.5. This method ranked peptides by the immunogenicity potential, taking into account NetMHCpan affinity, gene expression level, and mutant allele frequency. We then selected 153 predicted epitopes for functional validation, of which 33 were characterised positive by T-cell assay^63^. With only a small proportion of the shortlisted candidates found to be positive, the other existing models showed comparable levels of performance, PR-AUC ranging from 0.201 to 0.437 in our benchmarking study^7^.

Of the 153 tested peptides, we filtered out predicted HLA-A*02:01 non-binders and retained 9-10 amino acids peptides that were applicable to TRAP. This left 124 GBM peptides, of which 25 were characterised to be immunogenic and 99 non-immunogenic by T-cell assays. Of these, TRAP predicted 9/25 epitopes to be Positives and 84/99 non-epitopes to be Negatives (Figure 6E), yielding 0.75 accuracy with ROC-AUC of 0.684 (Figure 6F).

Identifying cancer neoepitopes is typically regarded as a ‘needle in a haystack’ problem in which an extremely small number of positives are sought from vastly imbalanced data. In this scenario, minimising the loss of true Positives from the candidate list was regarded to be more important than maximising accuracy. Therefore, we examined the confidence of predictions using the out-of-distribution detection module (Figure 6G) and found that many incorrectly predicted Negatives had low-confidence predictions (Figure 6H). By adding them to the candidate list, 24 more glioblastoma antigens were predicted positive, yielding 18/25 epitopes (Figure 6I) and ROC-AUC of 0.705 (Figure 6J).

Here, we applied TRAP for shortlisting glioblastoma cancer neoepitopes and showed that TRAP not only outperforms existing algorithms but also allow optimising candidates to minimise loss of likely epitopes.

## Discussion

In this study, we present TRAP (T-cell recognition potential of HLA-I presented peptides), a robust deep learning platform for predicting CD8+ T-cell epitopes from MHC-I presented pathogenic and self-peptides. To address the current limitations, we used peptide sequences only at contact positions to avoid HLA bias caused by capturing dominant HLA binding features at anchor positions. Second, we built separate models for pathogenic and self-peptides to address the out-of-distribution generalisation problem. Third, to overcome data limitations, we encoded peptide sequences using amino acid properties derived from protein transformer-based pre-trained language models (PLMs). This enabled more information about the physicochemical, electrostatic or biophysical properties of amino acids to be fed into the system. Fourth, we captured T-cell recognition motifs using a one-dimensional convolutional neural network (1D CNN) architecture. Fifth, we added the MHC binding rank score predicted by the most accurate algorithm (NetMHCpan) to provide the information about MHC binding. Lastly, we devised an out-of-distribution detection module to abstain from low-confidence predictions for peptides that are very different from those in the training dataset. By taking these novel approaches, TRAP offers a more robust prediction compared to other machine learning algorithms.

Another metric proposed to estimate immunogenicity from cancer neoepitope studies was dissimilarity to self. However, the dissimilarity to self could not be applied to pathogenic peptides because pathogens have highly heterologous sequences compared to the human proteome. Therefore, a novel metric termed RSAT (Relative similarity to autoantigens or tumour-associated antigens) was developed to estimate the immunogenicity of peptides from emerging pathogens. To compensate for the high dissimilarity between pathogenic and human proteomes, this metric compares the similarity of pathogenic peptides to the reference human proteome (‘healthy’) with respect to ‘immunogenic’ self-peptides such as autoantigens or tumour-associated antigens. This allowed the identification of pathogenic epitopes despite their sequence dissimilarity to the human proteome.

The application of TRAP was demonstrated by using it to identify cancer neoepitopes from glioblastoma patients. We showed that the out-of-distribution detection module is well-suited for the ‘needle in a haystack’ problem by identifying those predicted negative with low confidence. In another study, we used TRAP to investigate the immune escape potential of SARS-CoV-2 variants^64^. By combining TRAP with *in silico* mutagenesis, we evaluated the extent to which all possible theoretical single point mutations can give rise to variants of concern and be detrimental to T-cell immunity. Based on the groundwork of this work, using models like TRAP to systematically evaluate the impact of mutations on the emergence of deleterious pathogens will be of great interest. Emerging pathogens have posed a significant threat in recent years, and new variants and pathogens are expected to rise in the coming years^65^. It is therefore critical to surveil variants of concern and assess the immune escape potential of these variants. Furthermore, in order to accurately determine deleterious variants, it is essential to assess the extent to which models trained by other pathogens can be generalised and thus used for emerging pathogens. Thus, an iterative process of refining the training data, model architecture and validating predictions should be followed to mitigate the impact of another pandemic.

For the self-antigen model, the autoantigens, tumour-associated antigens and cancer neoepitopes were collected as epitopes, and benign HLA-I ligands expressed in thymus as non-epitopes. We assumed that HLA-I ligands (exclusively) expressed in the thymus are involved in the negative selection of T-cells, and thus there is no repertoire recognising these peptides. Notably, our novel approach of incorporating the HLA-I ligands allowed for a clearer separation of self-epitopes versus non-epitopes. When the data is limited, biological knowledge, such as negative thymic selection, can serve as a useful resource to bridge the data gap, and aid in the development of a more accurate classifier.

In benchmarking TRAP performance, we found that while Repitope performed well on pathogenic data, it was limited in extracting immunogenicity-related features from limited data and transferring them onto related species, resulting in poor performance on emerging pathogens. Our previous analysis revealed that Repitope was one of the models with skewed prediction for prevalent HLAs, with HLA-A*02:01-bound peptides having a higher immunogenicity score than non-HLA-A*02:01 bound peptides^7^. This suggested that the model considered A02:01 binding as a superior feature to T-cell recognition potential, skewing the prediction. While TRAP performed slightly lower on the pathogenic dataset, the HLA-generalised approach of TRAP mitigates the possibility of HLA bias. Therefore, end users should choose which model to pursue based on the peptides and HLA alleles of interest.

While these approaches improved the accuracy and robustness of the prediction, there still remains limitations. Foremost, while calibration methods could effectively detect incorrect prediction in 10-fold cross-validation, they failed on the cross-species dataset (Supplementary Figure 7G-H). We surmised this is because of “spurious semantic features” (i.e. features that have discriminative power in training data but not in test data) that were driving overly confident incorrect predictions. In a recent case study, Arora *et al*. benchmarked calibration methods on real-world challenge data and reported that models were over-confident on the OOD examples because of spurious semantic features and often produced accuracies close to random^45^. With spurious semantic shift being one of the remaining challenges in the NLP, further advances in deep learning and NLP will also facilitate improving the robustness of predictions on complex biological data.

Second, following our workflow, peptides may be abstained due to low-confidence predictions. For cancer peptides, TESLA^8^ that incorporates generic features for neoepitope prediction could be an alternative solution. The pathogenic peptides having low-confidence predictions can be directed to RSAT to estimate immunogenicity potential with respect to self-epitopes. However, because RSAT is only applicable to peptides with self-epitope counterparts, its coverage remains limited. Further characterisation of self-epitopes will expand RSAT coverage to embrace a broader range of pathogenic peptides.

Third, TRAP takes an HLA-generalised approach to avoid HLA bias, as the current dataset contains little intra-HLA variation in peptide immunogenicity. However, characterizing peptide immunogenicity on a broader range of HLAs may provide insight into the effect of HLA on overall immunogenicity. Moreover, because the specific interaction between peptide and T-cell receptor is MHC-restricted^66,67^, MHC-focused studies would be required to model specific TCR:pMHC recognition.

Fourth, T-cell recognition depends not only on peptide-MHC complex, but also on multiple other factors, such as cytokine microenvironment, co-stimulatory molecules *in vivo* and availability of TCR repertoires that are often highly stochastic and individualized^68^, adding complexity to T-cell activation and function^69^. While there are almost no record of these determinants^70^, studies highlighting the influence of microenvironment on T-cell response will be pivotal to recapitulating T-cell function for therapeutic applications.

In addition, we incorporated peptide sequence at the contact position to gauge on T-cell recognition motifs. However, some studies showed that position 1 (P1) of 9-mer peptide may play an important role in T-cell binding, as demonstrated by P1, 3, 4, 5, 9 being critical for MAGE A3 binding by a3a-engineered T-cells^71^. Although it is difficult to deconvolute the roles of different positions in MHC and T-cell binding, more studies may shed light on the role of P1.

Fifth, due to the lack of true Negative self-antigens, we retrieved benign HLA-I ligands exclusively expressed in thymus for their relevance to central tolerance. While there have been studies describing Tregs reacting to self-peptides present in the thymus^72,73^, given that the models are targeted for CD8+ T-cells, we rationed that majority of benign HLA-I ligands will be non-immunogenic. However, not all peptides expressed in thymus will be involved in negative selection, with the notion that positive selection-inducing self-peptides are displayed by cortical thymic epithelial cells (cTEC)^74^. Therefore, future studies characterizing HLA-I ligand expressed in medulla thymic epithelial cells (mTEC) will improve confidence as ‘non-immunogenic self-antigens’.

Sixth, it will be valuable to extend the work of peptide immunogenicity to investigate the ability of peptides to be recognised by *specific* T-cells. Here, the peptide immunogenicity was investigated at the organismal level i.e. whether the peptide can elicit a response from *any* T-cell. The limitation was largely due to the limited pool of available peptides characterised for their cognate TCRs, however, advances in screening methods will lead to availability of more comprehensive datasets in the future, thus enabling the development of more tailored immunogenicity models. It is becoming apparent that predicting specific interactions between TCR and cognate pMHCs is crucial for developing personalised therapies and tailoring vaccines or treatments to individuals’ TCR repertoire. Therefore, screening antigen-specific TCR against a larger pool of epitopes from various origins and pathologies will greatly aid in learning peptide features that allow interaction with specific TCRs.

Lastly, the current breadth of peptides characterised by T-cell assays is far from filling the full combinatorial peptide space, especially for CD4+ T-cell targets. Also, while this model is limited to peptide specific binding of CD8+ T-cells, other T-cells are specific for lipid or small molecules like metabolites. As such, high-throughput screening of immune targets and antigen-specific TCRs by the help of recent technological advancement will greatly foster the process of model development.

Despite current data and model constraints, the novel computational strategies allowed TRAP to outperform existing models in predicting CD8+ T-cell epitopes and provide more robust, accurate, and biologically meaningful candidates for functional validations. We believe that this platform will foster a better understanding of TCR:pMHC interaction and aid in basic, clinical and translational research for a wide range of therapeutic applications.

## Methods

### Data preparation

#### PeptideTcell data

Peptides that bind MHC-I molecules are typically restricted to 8-10 amino acids (aa) in lengths due to closed structure of peptide-binding groove. Given the limited number of 8aa peptides in databases, peptides of lengths 9-10aa characterized by T-cells were retrieved from IEDB. These included cancer neoepitopes, autoantigens or pathogenic peptides. The peptides without HLA allele or serotype annotations were removed. To ensure MHC binding, peptides were subjected to NetMHCpan 4.0 prediction and only those with rank ≤ 2.0 (i.e. predicted MHC binder) were retained. Peptides having contradictory immunogenicity annotations were categorized as ‘Positive’ and we only included Negatives that were characterised negative in ≥3 tests. This resulted 5,093 immunogenic (‘Positive’) and 6,628 non-immunogenic (‘Negative’) peptides in PeptideTcell data. The dataset include information about peptide sequence, binary immunogenicity, HLA allele, source antigen and organism. This data is the foundation for sequence pattern analysis, model development, RSAT validation.

#### Pathogenic data

For the analysis of sequence patterns and the development of deep learning-based models, datasets for pathogenic and self-antigens were prepared separately. For pathogenic datasets, we subset peptides originating from non-human species, resulting 4,000 Positive and 6,097 Negative pathogenic peptides.

#### Self-antigen data

For immunogenic self-peptides, autoantigens, tumor-associated antigens and cancer neoepitopes having 9-10 amino acids were collected from different databases. We gathered 162 epitopes from Cancer peptide database, 228 from dbPepNeo, 1506 from IEDB, 256 from McPAS-TCR, and 256 from NEPdb. For Negative self-antigens, we gathered HLA-I ligands expressed in thymus suggested to be involved in the negative selection of T-cells. 240 HLA-I ligands were collected from Adamopoulou *et al*., which were expressed in negatively selecting dendritic cells, 187 HLA-I ligands from Espinosa *et al*., which were expressed in thymus, and 10,840 benign HLA-I presented peptides from HLA Ligand Atlas, exclusively expressed in thymus. Of note, we did not include ‘Negatives’ from IEDB for self-antigen data because many were tested due to their association with tumour-associated antigens e.g. cancer/testis antigen 1, melanoma-associated antigen 9; yet, there was little evidence that these peptides were immunogenic. Therefore, only MHC-I peptides expressed in thymus were included. After pre-processing, removing duplicates and filtering for peptides with NetMHCpan rank ≤ 10, the self-antigen dataset was composed of 1,606 Positive and 10,915 Negative peptides. Notably, we expanded the MHC binding rank score range to include as many self-epitopes as possible for model training and validation.

#### Benchmarking data

The benchmarking analysis has been done in HLA-agnostic (i.e. peptide-based) or HLA-restrictive (i.e. peptide-HLA based) manner depending on the nature of different models. The TRAP makes prediction based on the peptide sequence and HLA binding rank score, which allows it to predict in both HLA-agnostic and restrictive manner. The HLA agnostic approach was applied on all peptides in the aforementioned pathogenic and self-antigen datasets. The HLA-restricted prediction was made against epitopes that were bound to 13 HLAs for which NetTepi could be performed, which are HLA-A*02:01, HLA-B*58:01, HLA-B*15:01, HLA-B*35:01, HLA-B*07:02, HLA-A*01:01, HLA-A*03:01, HLA-A*11:01, HLA-A*24:02, HLA-A*26:01, HLA-B*27:05, HLA-B*39:01, HLA-B*40:01. The list of models conducted in HLA-restricted manner are iPred, PRIME, NetTepi, IEDB and DeepImmuno. Specifically for DeepImmuno, an additional filter was applied to exclude peptides that were bound to HLAs that DeepImmuno could not process.

#### DeepImmuno data

DeepImmuno training data was retrieved from GitHub repository, and used for evaluating the DeepImmuno performance. https://github.com/frankligy/DeepImmuno

### 10-fold CV and cross-species comparison

The peptideTcell data was divided randomly or in cross-species manner: i) 90% train vs. 10% test random split (i.e. representation of 10-fold cross-validation), ii) Non-SARS-CoV-2 (Non-SARS-2) train vs. SARS-2 test, and iii) Non-vaccinia virus (Non-VACV) train vs. VACV test.

To compare the performance of the XGBOOST classifier on random vs. cross-species datasets, amino acids at contact positions were first represented by their physicochemical properties using “aaDescriptors” function in R Peptides v2.4.4 package. The amino acid descriptors included kideraFactor, zScales, tScales, vhseScales, protFP, stScales, blosumIndices, mswhimScores, crucianiProperties and fasgaiVectors, which described properties such as polarity, electronic properties, hydrophobicity, α-helix/bend preference, β-sheet, bulkiness/size of side-chains, hydrogen-bonding, isoelectric point and structural topology^75^. In addition to amino acid-level embedding, the peptide-wide property was added by averaging these aaDescriptors across all positions for each peptide. The embeddings from random split and cross-species datasets were used to generate the XGBOOST classifier using “XGBClassifier” function in python xgboost v0.90 package. The set of hyperparameters, such as alpha, gamma, max_depth and colsample_bytree, were optimized by grid search for each dataset. The models were trained using training datasets: 90% train, Non-SARS-2 and Non-VACV peptides by 10-fold cross-validations. The trained model were tested on 10% test, SARS-2 and VACV peptides, respectively as representatives of 10-fold CV and cross-species comparisons.

To analyse sequence homology between training vs. test datasets, differential position specific scoring matrices (dPSSMs) were generated for each training and test datasets. The probability frequency of each amino acids in each position was represented by position specific scoring matrices using “consensusMatrix” function from R Biostrings v2.56.0 package. The PSSMs from Positive and Negative peptides were standardised by centre and scaling, and differential PSSMs were generated by subtracting the two. To estimate the discriminative power of the dPSSM scores, we generated dPSSMs using training data and used the matrices to score respective test peptides for their immunogenicity potential.

### Immunogenicity positivity score

The positivity score was computed by taking three factors into account: 1) the number of experiments conducted, 2) the percentage tested positive and 3) the number of cognate TCRs if available, using the following equations.

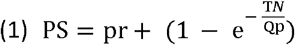

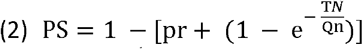

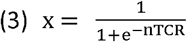

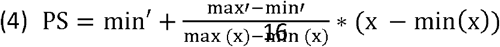

(1) computes positivity score (PS) for Positives peptides that do not have TCR information available, where pr = % responded positive, TN = total number of tests conducted and Qp = the number of tests designated as a minimum number of tests required to support positivity (Qp = 3). (2) computes PS for Negative peptides, where Qn = number of tests designated as a minimum number of tests required to support negativity (Qn = 5). (3) and (4) compute for Positives with cognate TCR information. (3) is a sigmoid function that translates the number of cognate TCR (nTCR) to values in a logistic growth curve and (4) scales the distribution to align with scores from (1), where min’ equals mean of the distribution (~1.28 equivalent to 1 test and 1 responded positive) and max’ = 2.0. Due to limited pool of peptides with cognate TCRs, the majority of peptides’ positivity score was computed by translating the number of experiments conducted and % tested positive, and the number of cognate TCRs was incorporated to add greater weights to positivity (all positivity scores with cognate TCR had values > 1). The positivity scores were computed for each entry for each peptide-HLA pair, and ranged from 0 to 2.3, with Negatives ranging from 0-1 and Positives ranging from 1-2.3 (Supplementary Figure 4D).

### Intra- and inter-HLA variability

The HLA effect was computed on 2,349 peptides having entries from multiple HLA alleles. One-way ANOVA was conducted for the effect of HLA on the positivity score for each peptide-HLA pair using ‘aov’ function from R stats v4.0.5 package. The distributions show mean squared from summary output.

### Differential sequence patterns

The n-grams (i.e. contiguous sequence of n-amino acids) were generated using ‘ngram’ function from R ngram v3.1.0 package. The number of peptides containing the n-grams was counted for Positive and Negative respectively and normalised for the total number of Positive and Negative peptides, respectively. Then, we computed ratio_n-gram_ = normalized # of Positive peptides containing the n-gram / normalized # of Negative peptides containing the n-gram, and shortlist top differential n-grams by the ratio score. Similarly, we generated all possible combinations of position-specific k-mer motifs (i.e. contiguous or non-contiguous sequence of k amino acids restricted to peptides of same lengths), where e.g. .M.W. denotes MW pattern at P2 and P4 of 5 amino acid peptide. We computed ratio _positional k-mer_ = normalized # of Positive peptides containing the positional k-mer / normalized # of Negative peptides containing the positional k-mer to shortlisted top differential position-specific k-mer motifs. For visualization, patterns were categorized by their normalized ratios, where ‘lows’ have ¼ < ratio < 4, ‘high in pos’ have ratio ≥ 4, ‘high in neg’ have ratio ≤ ¼, and ‘onlys’ have motifs in either positive or negative sets, and top differential patterns from ‘high’ or ‘only’ categories were visualized by barplot. To identify shared enriched patterns between pathogenic and self-antigens, only n-grams or positional k-mer motifs having ratio ≥ 3 or ≥ 3 positive peptides (for onlys) were pre-selected for comparison. The pairwise sequence similarity between peptides was computed using ‘pairwiseAlignment’ function in Biostrings v2.56.0 package^76^, using BLOSUM62 substitution matrix and default parameters. For clusters of highly similar peptides, the peptides having alignment scores ≥22 with ≥3 other peptides were visualized into network graph using R ggnetwork v0.5.10 package. The toolkits for generating the sequence patterns are deposited in: https://github.com/ChloeHJ/diffSeqPatterns.

### Deep learning models for pathogenic and self-epitope prediction

The peptide sequences at contact positions, i.e. P3-P8 of 9aa and P3-P9 of 10aa peptides, were encoded either by one-hot-encoding having dimension [m, 7, 21], amino acid descriptors [m, 7, 49] or amino acid embeddings from protein transformer-based pretrained language models (PLMs) models [m, 7, 1024], where m represents the number of peptides. The amino acid descriptors included kideraFactors, tScales, protFP, BLOSUM, stScales and MSWHIM captured by “aaDescriptors” function from R Peptides v2.4.4 package, and Atchley factors from “AAMetric.Atchley” function in R HDMD v1.2 package. In addition, amino acids were embedded using five protein transformer-based PLMs, including prot_t5_xl_uniref50, prot_bert, prot_bert_bfd, prot_t5_xl_bfd, prot_xlnet from Rostlab using Tokenizer and EncoderModel functions from python transformers v4.19.0 package. These models are based on T5 or BERT and were pre-trained on a large corpus of protein sentences, e.g. UniRef50, a dataset consisting of 45 million protein sequences, in a semi-supervised fashion. Further details about transformer-based PLMs can be found in https://huggingface.com/Rostlab.

To account for peptides of varying lengths, the peptides coming from 9aa peptides were padded (i.e. adding non-relevant number to the borders of the matrix) either in the front (i.e. pre-padding) or at the back (i.e. post-padding). First, we computed all possible n-grams from 6- and 7-mer peptides, and analysed if the same n-grams were present in both 6- and 7-mers. We found that many n-grams were present in both 6- and 7-mers. We then aligned by their respective positions, and observed that many 3-grams located in P3-P5 of 9aa peptides were positioned at P4-P6 of 10aa peptides, indicating padding in front of 6-mer peptide (i.e. pre-padding) would align with hotspots in 7-mers.

The classification accuracies of different deep learning architectures were compared between simple dense layer (i.e. classification), biRNN, biLSTM, 1D CNN, 2D CNN and transformer models. The biRNN model contained two biRNN layers each having 512 units, followed by a dense layer of 256 units and a classification layer with dropouts in-between. The biLSTM model had the same structure as the biRNN model, but LSTM cell is used instead of the RNN cell. 2D CNN model had two Conv2D layers, with filters = 16 and 32 respectively, followed by MaxPool2D, Flatten, a dense layer of 256 units anda classification layer with dropouts in-between. The transformer model for pathogenic and self-antigen data had 10 and 2 attention heads respectively and 32 hidden layers in a feed forward network, followed by GlobalAveragePooling1D, dense layer with 128 units and classification, with dropouts in-between. The 1D CNN had kernel sizes 1, 3, 5 and 7, each max pooled and concatenated to a layer. In parallel, -log transformed NetMHCpan rank and hydrophobicity (i.e. the proportion of A, V, L, M, W) have been added as a MLP layer. 1D CNN and MLP layers were concatenated, and put into a dense layer of 256 units followed by classification. The hyperparameters of the final 1D CNN models were optimized by grid search. The final hyperparameters for pathogenic model are: learning rate = 1e-05, weight decay = 1e-06, dropout rate = 0.1, batch size = 50, dense layer node = 2000 and dense layer node = 256, giving ROC-AUC of 0.764 by 10-fold cross-validation. The final hyperparameters for self-antigen model are: learning rate = 0.001, weight decay = 0.01, dropout rate = 0.2, batch size = 100, MLP dense = 1500 and dense layer node = 512, giving ROC-AUC of 0.943 by 10-fold cross-validation. All deep learning architectures are implemented using python TensorFlow v2.8.0 package.

### Out of distribution (OOD) detection

Autoencoder is a type of unsupervised neural network that have a smaller number of neurons in the hidden layers than the input layer. This allows the architecture to extract essential information from the input layer, preserve in lower-dimension, and employ to reconstruct output. We used the difference between the input and reconstructed output (i.e. reconstruction loss) as a metric for anomaly detection, as autoencoders cannot effectively reconstruct patterns not learnt from training data^56^.

For anomaly detection, peptides encoded by prot_t5_xl_uniref50 transformer-based PLM were applied onto 2D and 3D autoencoders, variational autoencoder and denoising autoencoder models, using ‘AutoEncoder’ function from python pyod v1.0.1 package. The autoencoder models were trained using 90% random train and non-SARS-CoV-2 data, and were used to predict 10% test and SARS-CoV-2 data respectively. Then, we computed reconstruction loss between original and predicted test values.

For calibration methods, the 1D CNN model was trained using softmax activation function. We then computed softmax probability (MaxProb) and temperature-scaled softmax probability (T=2) for each test peptide. The MaxProb is the maximum softmax probability between class 0 (Negative) and 1 (Positive). The temperature scaling softens the softmax probability with T > 1, making the network slightly less confident, reflecting the true probabilities^77^. For Ensemble, we used 10-folds of the training dataset to generate 10 different models. These models were then used to predict test data points, producing prediction scores for each. We combined MaxProb and found the average MaxProb of ten models. For Monte Carlo dropout, Monte Carlo models were reiterated 100 times with stochastic dropout of 0.6. The MaxProb was generated from each, and averaged over 100 scores. The final OOD classifier predicts correct vs. incorrect predictions by using the MaxProb and average MaxProb from the Ensemble as inputs to the decision tree classifier. The classifier was built using the ‘DecisionTreeClassifier’ function from python sklearn v1.0.2 package.

### Relative similarity to autoantigens or tumour-associated antigens (RSAT)

To compute RSAT, a total of 5023 unique cancer neoepitopes, autoantigens, tumour-associated antigens and other self-epitopes were retrieved from IEDB, dbPepNeo, NEPdb, McPAS-TCR and tumour antigenic peptide database. First, only pathogenic peptides having comparable self-epitope counterparts are retrained by computing Match score^78^ (5) between pathogenic peptides and self-epitopes, where BL represents the global-local alignment score using BLOSUM62 matrix, p represents pathogenic peptide and se self-epitopes. Only pathogenic peptides having a match score ≥ 0.6 are retained to compute RSAT.

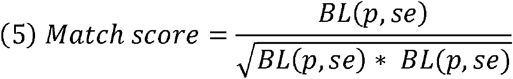

For pathogenic peptides having comparable self-epitope counterparts, RSAT was computed (6). First, we computed the maximum global-local alignment score between pathogenic peptides and AATs (a set of autoimmunity, allergy tor tumour-associated antigens) using the BLOSUM62 substitution matrix. Second, we computed the maximum global alignment score between pathogenic peptide and AAT’s best counterpart in human proteome. Third, we computed the ratio between alignment scores from self-epitope and human proteome counterpart. In the equation (6) below, p = pathogenic peptide, AAT = autoimmunity, allergy tor tumour-associated antigens, hp = human proteome.

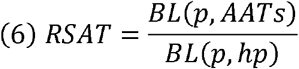

### Application of TRAP to shortlisting glioblastoma neoantigens

In previous in-house study^63^, tumours of four HLA-A2 glioblastoma patients were sequenced and cancer neoepitopes were shortlisted using an in-house version of MuPeXI^62^ named TUNAPASTA v0.5. We then selected 153 predicted neoepitopes for functional T-cell assays, 33 of which were characterised Positive. From these 153 tested peptides, we filtered out predicted HLA*02:01 non-binders and retained 9-10 amino acid peptides that were applicable to TRAP. This left 124 GBM peptides, 25 of which were Positives and 99 Negatives. We used a pre-trained self-antigen TRAP model on this dataset to predict the immunogenicity of GBM peptides as well as the confidence of prediction by the OOD detection module. Because identifying cancer neoepitopes is thought to be a ‘needle in a haystack’ problem, we identified predicted Negatives with low-confidence prediction, and added 24 more candidates for validation, resulting in a ROC-AUC of 0.705.

## Author contributions

CL and HK conceived the study. CL designed and conducted all computational analysis with inputs from AA, JH, MJ, PB, MP, RF, AS and HK. CL developed deep learning model with inputs from JH and MJ. PB and CL conducted benchmarking. CL wrote the manuscript. JH, PB, MJ, MP, RF, AA, AS and HK commented and edited the manuscript. AS and HK supervised the study. All authors contributed to the article and approved the submitted version.

## Acknowledgements

This work has been supported by Medical Research Council. HK is funded by MRC Human Immunology Unit core funding. CL is funded by UK National Institute of Health Research (NIHR). AS is funded by a Wellcome Investigator Award (219523/Z/19/Z), the UK Medical Research Council, NIHR, awards from Bristol-Myers Squibb and UCB. The views expressed are those of the author(s) and not necessarily those of the NHS, the NIHR, or the Department of Health, UK.

## Supplementary Figure Legends

**Supplemetnary Figure 1. Screnshot of TRAP web application.** TRAP has been developed as a user-friendly web application, where users can enter their peptide list and select the model of interest (pathogenic or self-antigen), and the application will compute the prediction scores along with its confidence. The users can export figure and/or table for their downstream applications.

**Supplementary Figure 2. Cross-species variation and HLA-bias.** A. Statistics of peptides in pathogenic data by their species of origin. B. Distribution of MHC binding rank scores (-log2 transformed) predicted by NetMHCpan 4.0 on pathogenic peptides. C. Distribution of hydrophobicity (i.e. the proportion of hydrophobic amino acids, A, V, L, M and W) of the pathogenic peptides. D. Distribution of differential position specific scoring matrix (dPSSM) score. The dPSSMs were first generated by using training datasets i.e. 90% random data, non-SARS2, non-VACV peptides and were used to predict dPPSM scores on the test dataset i.e. 10% random data, SARS2 and VACV peptides. E. Statistics of peptides in DeepImmuno training dataset. F. Performance of DeepImmuno. The ROC curve was reproduced as published in the original publication by 10-fold cross-validations. G. ROC curve showing DeepImmuno performance on randomly sampled 3303 peptide-HLA pairs (i.e. equivalent number of peptide-HLA pairs as HLA*02:01) to demonstrate that reduced accuracy of single-HLA predictions is not due to lower number peptides in the datasets. H. Statistics on the number of peptides per HLA after down-sampling to balance the number of epitopes and non-epitopes.

**Supplementary Figure 3. Effect of anchor and contact positions on peptide immunogenicity.** A-B. t-SNE embedding of peptides represented by peptide-wide descriptors and amino acids at anchor positions, coloured by cognate TCRs for 9aa peptides (A) and 10aa peptides (B). C. t-SNE embeddings of peptides represented by peptide-wide descriptors and position-specific amino acid descriptors, coloured by supertype, immunogenicity, MHC binding and source organism of each peptide. The t-SNEs contain amino acid residues at different positions, in which left plots contain amino acids at all positions, middle plots contain amino acids at contact positions and right plots contain amino acids at anchor positions.

**Supplementary Figure 4. Intra vs. Inter-HLA variability.** A-C. Convert binary to quantitative positivity score. The positivity score is computed using the number of tests conducted (A), the percentage that responded positive (B) and the number of cognate TCRs (C). D. Distribution of positivity score. In essence, for positive peptides, the higher the positivity score, more likely the peptide is to be immunogenic. For negative peptides, the lower the score, more likely the peptide to be non-immunogenic. E. Example of peptides having higher intra-HLA variation than inter-HLA variation. F. Sources of intra-HLA variation. The causes of intra-HLA variation include biological and technical factors, such as different assays, effector population and antigen-presenting cells in characterizing the immunogenicity of peptides bound on the same HLA. ‘-‘ means information is not available. For peptide-HLAs without supporting information, positivity scores were denoted as binary values.

**Supplementary Figure 5. Pathogenic and self-antigen datasets.** A. Statistics on the number of peptides in the pathogenic dataset. B. Statistics on the number of peptides in the self-antigen dataset. C. Statistics on the number of self-epitopes and HLA-I ligands expressed in thymus, retrieved from publications and databases.

**Supplementary Figure 6. Sequence patterns discriminating epitopes versus non-epitopes.** A-B. Sequence logo of differential amino acid usage in contact positions of 9aa and 10aa peptides in pathogenic (A) and self-peptides (B). C. Top n-grams enriched (green) or depleted (orange) in epitopes for pathogenic (top) or self-antigens (bottom). D. Example of top n-grams found in both pathogenic and self-epitopes. Shown is the number of peptides containing the n-grams. E. Normalized ratio of shared position-specific k-mer motifs found from 9aa peptides enriched in both pathogenic and self-epitopes. F-G. Example of top epitope-enriched position-specific k-mer motifs found in both pathogenic and self-peptides for 9aa (F) and 10aa (G). Shown is the number of peptides containing the position-specific k-mer motifs. H-I. Clusters of peptides with high sequence similarity in contact positions, demonstrated by pairwise global alignment scores. Heatmap showing 9aa (H) and 10aa (I) pathogenic peptides having global alignment score ≥ 27 with ≥ 3 other peptides.

**Supplementary Figure 7. OOD detection.** A-B. Anomaly detection using autoencoders to discriminate correctly or incorrectly predicted pathogenic (A) and self- (B) peptides. Autoencoders included are 2-dimensional autoencoder (2D AE), 3-dimensional autoencoder (3D AE), variational autoencoder and denoising autoencoder. Statistical significance by p-values from Student’s t-test. ns: non-significant. We observed no significant difference in reconstruction loss between correctly and incorrectly predicted peptides, implying that the autoencoder-based methods cannot effectively identify OOD inputs. We speculated that unsupervised anomaly detection methods, such as reconstruction loss from autoencoders, were ineffective because ‘species’ was not the primary driver of peptide clustering at contact positions, which was often driven by the cognate TCRs to which they were bound. Likewise, density-based or distance-based methods that use sequence-representation similarity would not be able to detect OODs. C-F. Anomaly detection using autoencoders – 2D-AE (C), 3D AE (D), variational autoencoder (E) and denoising autoencoders (F) for discriminating correctly vs. incorrectly predicted pathogenic (top) and self- (bottom) peptides. For SARS-2 peptides, non-SARS-2 peptides were used to train autoencoders to predict correctly vs. incorrectly predicted SARS-2 peptides. Statistical significance by p-values from Student’s t-test. ns: non-significant. G-H Distribution of maximum softmax probability (MaxProb) (G) and average maximum softmax probability (average MaxProb) from 10 ensembled models (H) for randomly divided data (resemblance of 10-fold cross-validation) and cross-species data (i.e. model trained using non-SARS-CoV-2 peptides to predict SARS-CoV-2 peptides). I. MaxProb (left) and average MaxProb of 10 ensembled models (right) for glioblastoma cancer peptides predicted to be immunogenic using the TRAP self-antigen model.

## Supplementary Tables

Supplementary Table 1. Pathogenic dataset

Supplementary Table 2. Self-antigen dataset

Supplementary Table 3. Pathogenic TRAP model hyperparameter optimisation

Supplementary Table 4. Self-antigen TRAP model hyperparameter optimisation

Supplementary Table 5. Pathogenic sequence patterns

Supplementary Table 6. Self-antigen sequence patterns

Supplementary Table 7. Pathogenic OOD model calibration metrics

Supplementary Table 8. Self-antigen OOD model calibration metrics

Supplementary Table 9. GBM data

