## Supplemental Figures for "A robust deep learning platform to predict CD8+ T-cell epitopes"

#### Supplementary Figures

#### TRAP: Deep learning platform for CD8+ T-cell epitope prediction

Self-antigen model

Drag and Drop or Select Files

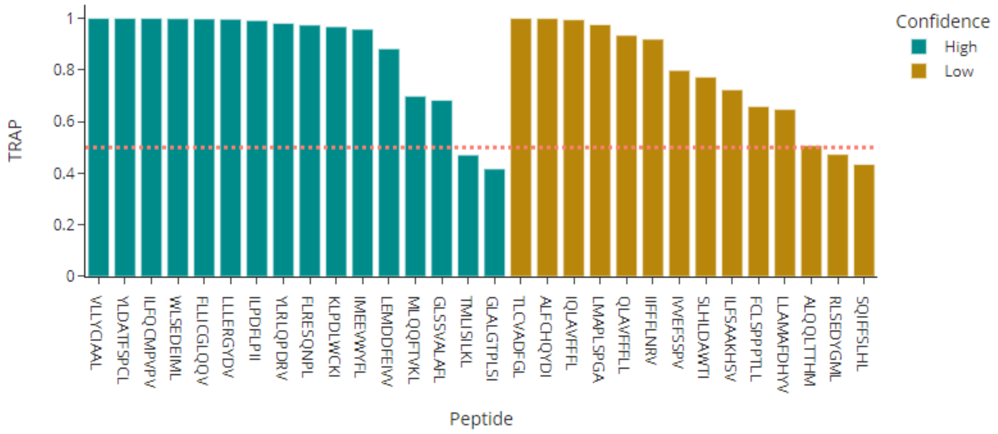


gbm_test_data.csv

EXPORT

| Peptide | ContactPosition | nlog2Rank | TRAP | MaxProb | Ensemble | Confidence |
|---|---|---|---|---|---|---|
| VLLYCIAAL | LYCIAA | -0.614 | 0.99988 | 7.361 | 5.878 | High |
| TLCVADFGL | CVADFG | -0.404 | 0.99987 | 5.689 | 5.076 | Low |
| YLDATFSPCL | DATFSPC | -0.019 | 0.99985 | 6.741 | 6.594 | High |
| ILFQCMPVPV | FQCMPVP | -0.568 | 0.99981 | 7.1 | 6.842 | High |
| ALFCHQYDI | FCHQYD | 0.199 | 0.99947 | 3.789 | 2.413 | Low |
| WLSEDEIML | SEDEIM | 4.658 | 0.99886 | 5.464 | 5.77 | High |
| FLLICGLQQV | LICGLQQ | 0.907 | 0.99786 | 5.972 | 6.011 | High |
| LLLERGYDV | LERGYD | 2.816 | 0.99673 | 1.681 | 1.921 | High |
| IQLAVFFFL | LAVFFF | -0.457 | 0.99444 | 0.128 | 1.588 | Low |
| ILPDFLPII | PDFLPI | 4.216 | 0.99175 | 2.844 | 2.743 | High |
| YLRLQPDRV | RLQPDR | 0.44 | 0.98089 | 3.361 | 1.266 | High |

**Supplemetnary Figure 1. Screnshot of TRAP web application.** TRAP has been developed as a user-friendly web application, where users can enter their peptide list and select the model of interest (pathogenic or self-antigen), and the application will compute the prediction scores along with its confidence. The users can export figure and/or table for their downstream applications.

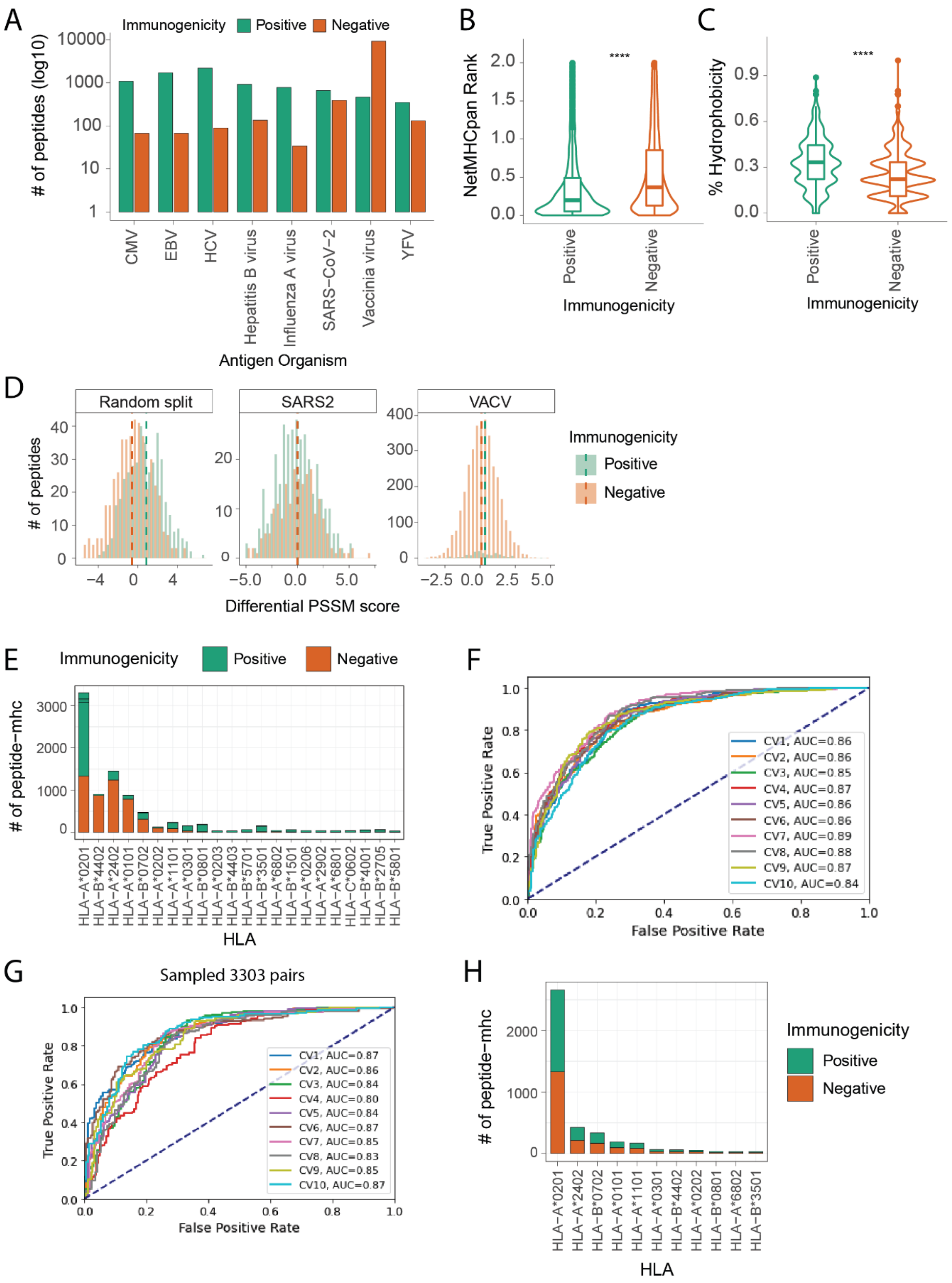


**Supplementary Figure 2. Cross-species variation and HLA-bias.** A. Statistics of peptides in pathogenic data by their species of origin. B. Distribution of MHC binding rank scores (-log2 transformed) predicted by NetMHCpan 4.0 on pathogenic peptides. C. Distribution of hydrophobicity (i.e. the proportion of hydrophobic amino acids, A, V, L, M and W) of the pathogenic peptides. D. Distribution of differential position specific scoring matrix (dPSSM) score. The dPSSMs were first generated by using training datasets i.e. 90% random data, non-

SARS2, non-VACV peptides and were used to predict dPPSM scores on the test dataset i.e. 10% random data, SARS2 and VACV peptides. E. Statistics of peptides in DeepImmuno training dataset. F. Performance of DeepImmuno. The ROC curve was reproduced as published in the original publication by 10-fold cross-validations. G. ROC curve showing DeepImmuno performance on randomly sampled 3303 peptide-HLA pairs (i.e. equivalent number of peptide-HLA pairs as HLA*02:01) to demonstrate that reduced accuracy of single-HLA predictions is not due to lower number peptides in the datasets. H. Statistics on the number of peptides per HLA after down-sampling to balance the number of epitopes and non-epitopes.

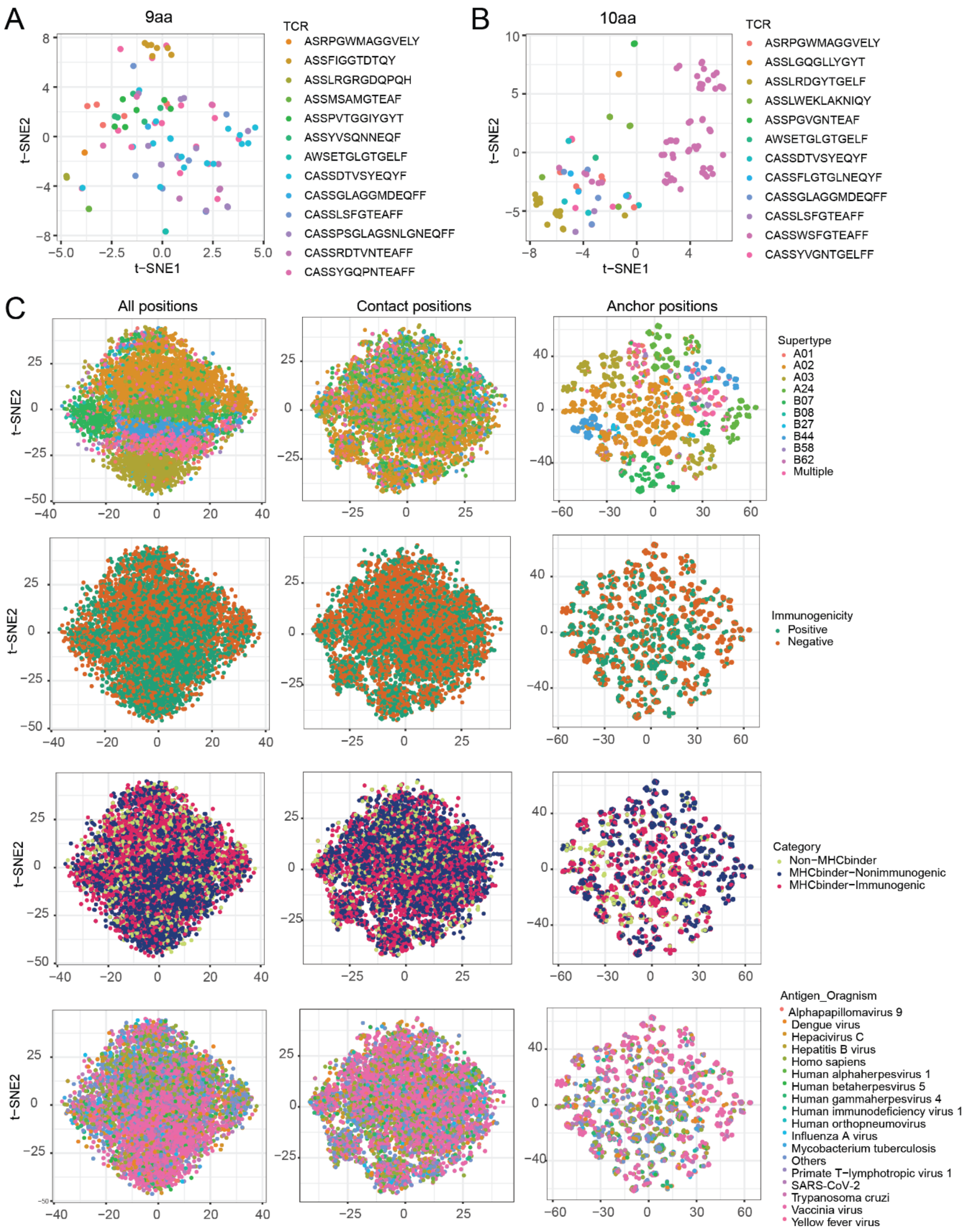


**Supplementary Figure 3. Effect of anchor and contact positions on peptide immunogenicity.** A-B. t-SNE embedding of peptides represented by peptide-wide descriptors and amino acids at anchor positions, coloured by cognate TCRs for 9aa peptides (A) and 10aa peptides (B). C. t-SNE embeddings of peptides represented by peptide-wide descriptors and position-specific amino acid descriptors, coloured by supertype, immunogenicity, MHC binding and source organism of each peptide. The t-SNEs contain amino acid residues at different positions, in which left plots contain amino acids at all positions, middle plots contain amino acids at contact positions and right plots contain amino acids at anchor positions.

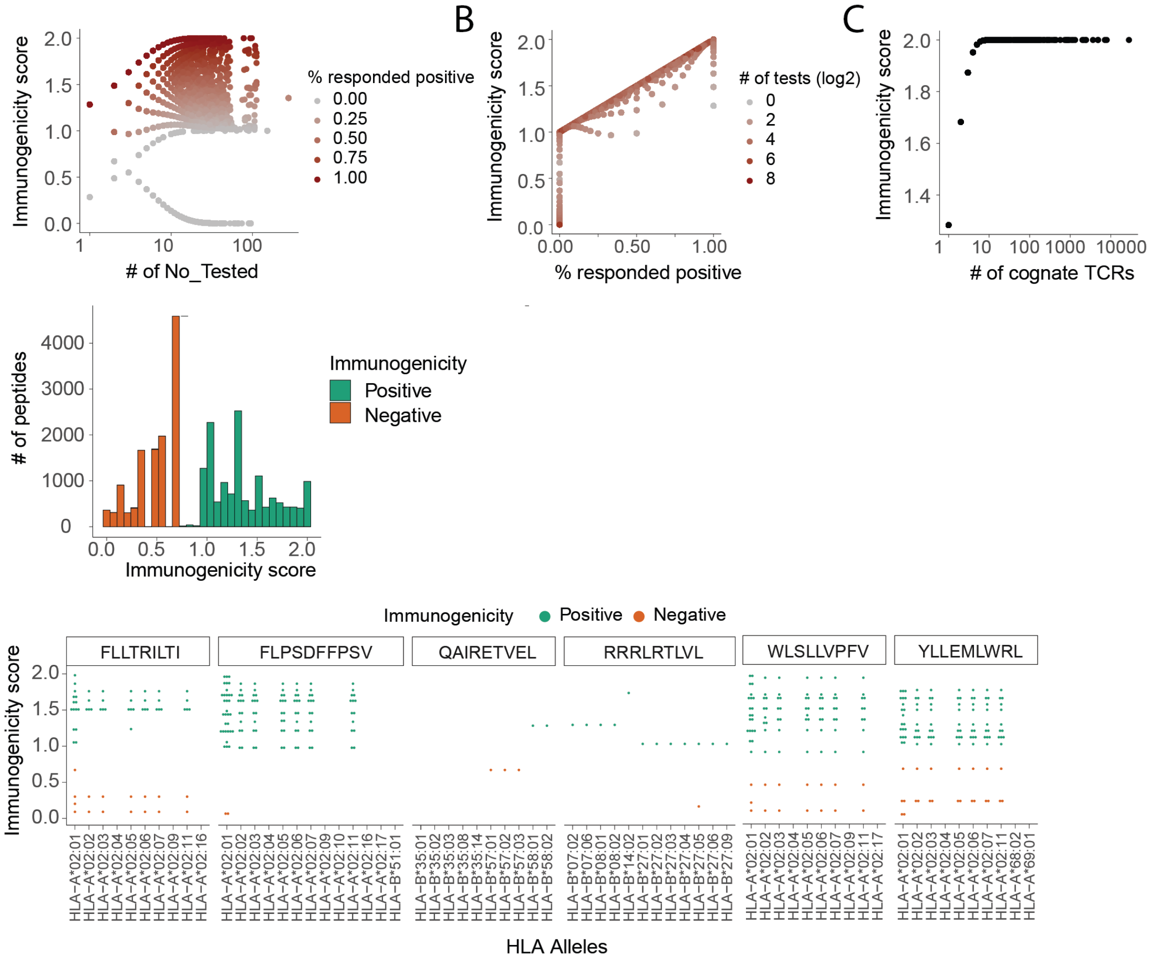


F

| Peptide | Immunogenicity | MHC Restriction | Antigen Organism | # Tested | # Responded | Response frequency | In vitro stimulation | Assay | Assay group | Effector cell culture condition | Antigen presenting cells |
|---|---|---|---|---|---|---|---|---|---|---|---|
| GILGFVFTL | Positive | HLA-A*02:01 | Influenza A virus | | | | Restimulation in vitro | 51 chromium | cytotoxicity | Cell Line / Clone | B cell |
| GILGFVFTL | Positive | HLA-A*02:01 | Influenza A virus | | | | Restimulation in vitro | ELISPOT | IFNγ release | Direct Ex Vivo | PBMC |
| GILGFVFTL | Positive | HLA-A*02:01 | Influenza A virus | | | | Restimulation in vitro | multimer/ Tetramer | qualitative binding | Short Term Restimulated | |
| GILGFVFTL | Positive | HLA-A*02:01 | Influenza A virus | 6 | 6 | 100 | | ICS | CXCL9/MIG release | Direct Ex Vivo | PBMC |
| GILGFVFTL | Positive | HLA-A*02:01 | Influenza A virus | 5 | 2 | 40 | | ELISPOT | IFNγ release | Direct Ex Vivo | PBMC |
| GILGFVFTL | Positive | HLA-A*02:01 | Influenza A virus | 1 | 1 | 100 | Restimulation in vitro | 51 chromium | cytotoxicity | Cell Line / Clone | B cell |

**Supplementary Figure 4. Intra vs. Inter-HLA variability.** A-C. Convert binary to quantitative positivity score. The positivity score is computed using the number of tests conducted (A), the percentage that responded positive (B) and the number of cognate TCRs (C). D. Distribution of positivity score. In essence, for positive peptides, the higher the positivity score, more likely the peptide is to be immunogenic. For negative peptides, the lower the score, more likely the peptide to be non-immunogenic. E. Example of peptides having higher intra-HLA variation than inter-HLA variation. F. Sources of intra-HLA variation. The causes of intra-HLA variation include biological and technical factors, such as different assays, effector population and antigen-presenting cells in characterizing the immunogenicity of peptides bound on the same HLA. '-' means information is not available. For peptide-HLAs without supporting information, positivity scores were denoted as binary values.

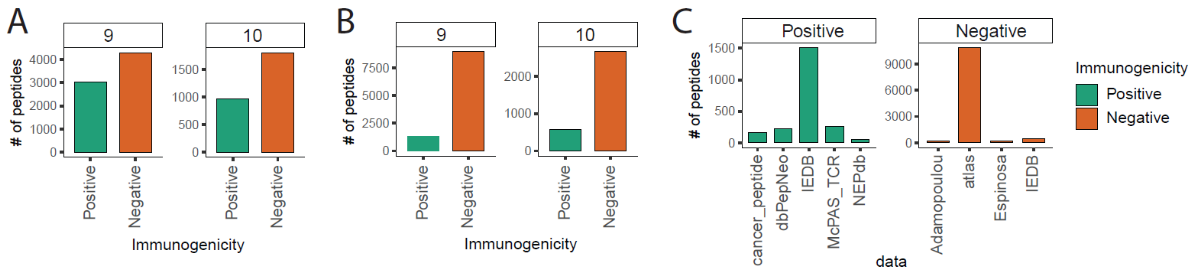


**Supplementary Figure 5. Pathogenic and self-antigen datasets.** A. Statistics on the number of peptides in the pathogenic dataset. B. Statistics on the number of peptides in the self-antigen dataset. C. Statistics on the number of self-epitopes and HLA-I ligands expressed in thymus, retrieved from publications and databases.

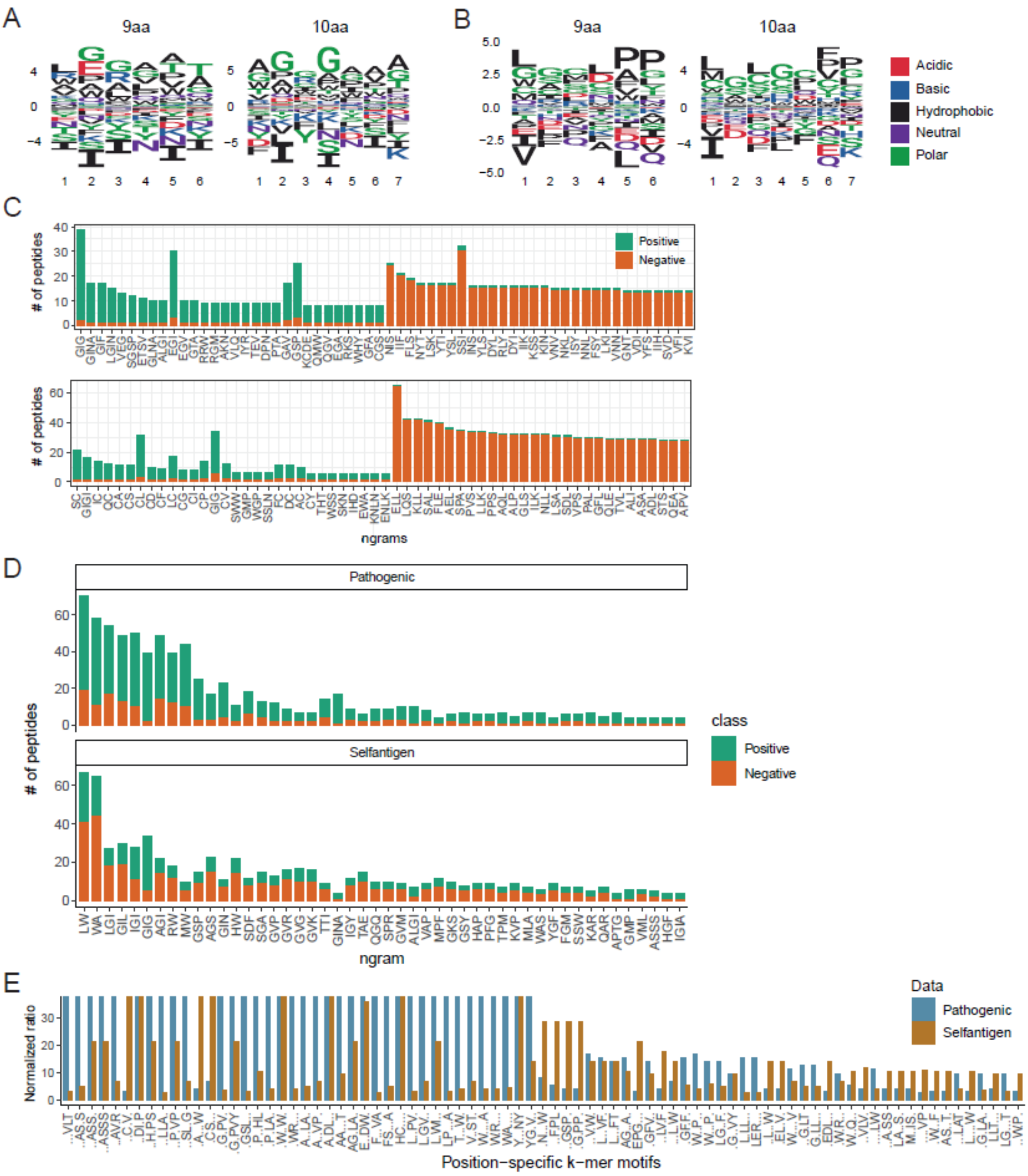
A
9aa
10aa
B
9aa
10aa
Acidic
Basic
Hydrophobic
Neutral
Polar
C
### of peptides
Positive
Negative
ngrams
D
Pathogenic
Selfantigen
### of peptides
class
Positive
Negative
ngram
E
Normalized ratio
Data
Pathogenic
Selfantigen
Position−specific k−mer motifs

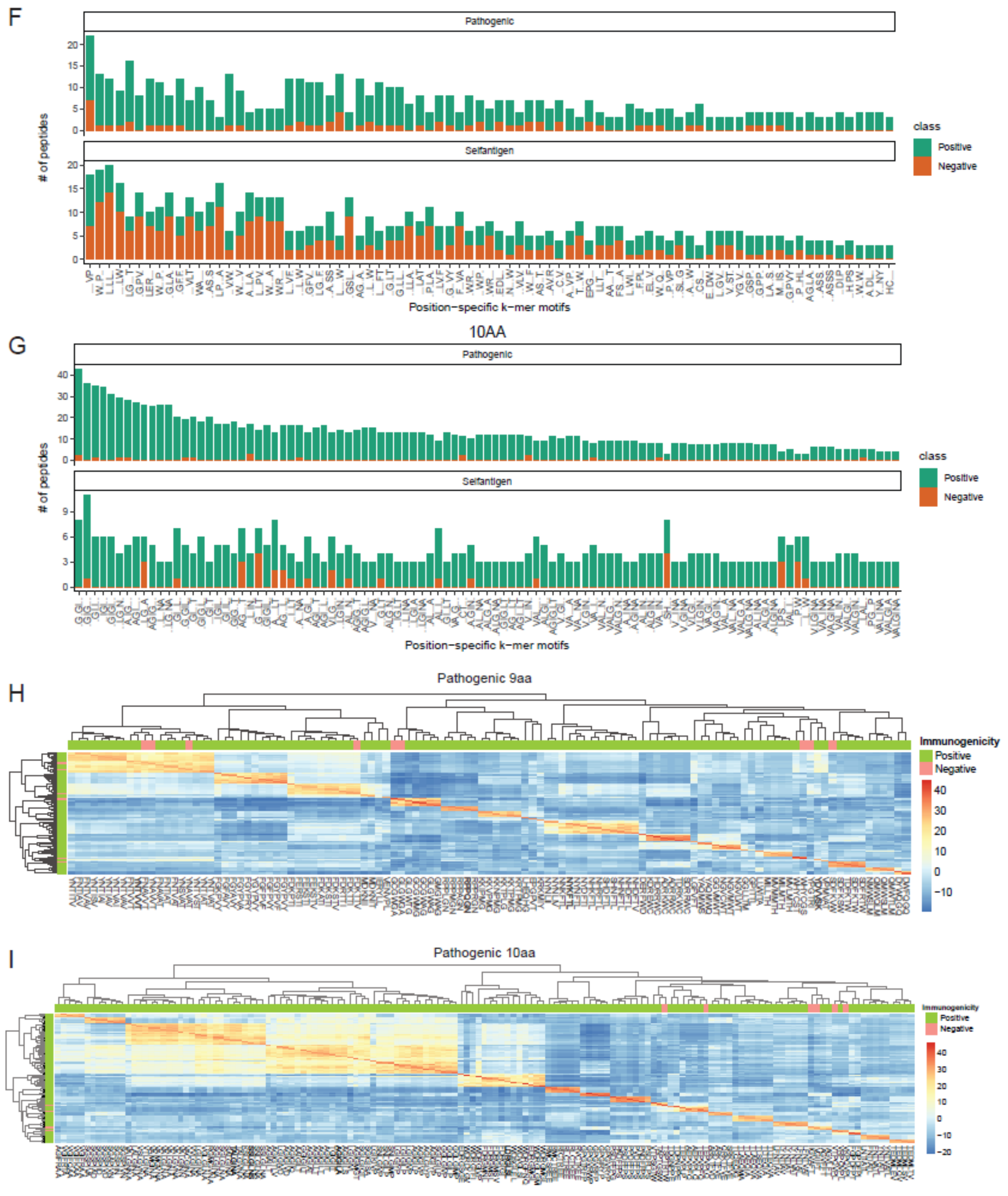


**Supplementary Figure 6. Sequence patterns discriminating epitopes versus non-epitopes.** A-B. Sequence logo of differential amino acid usage in contact positions of 9aa and 10aa peptides in pathogenic (A) and self-peptides (B). C. Top n-grams enriched (green) or depleted (orange) in epitopes for pathogenic (top) or self-antigens (bottom). D. Example of top n-grams found in both pathogenic and self-epitopes. Shown is the number of peptides containing the n-grams. E. Normalized ratio of shared position-specific k-mer motifs found from 9aa peptides enriched in both pathogenic and self-epitopes. F-G. Example of top epitope-enriched position-specific k-mer motifs found in both pathogenic and self-peptides for 9aa (F) and 10aa (G). Shown is the number of peptides containing the position-specific k-mer motifs. H-I. Clusters of peptides with high sequence similarity in contact positions, demonstrated by pairwise global alignment scores. Heatmap showing 9aa (H) and 10aa (I) pathogenic peptides having global alignment score ≥ 27 with ≥ 3 other peptides.

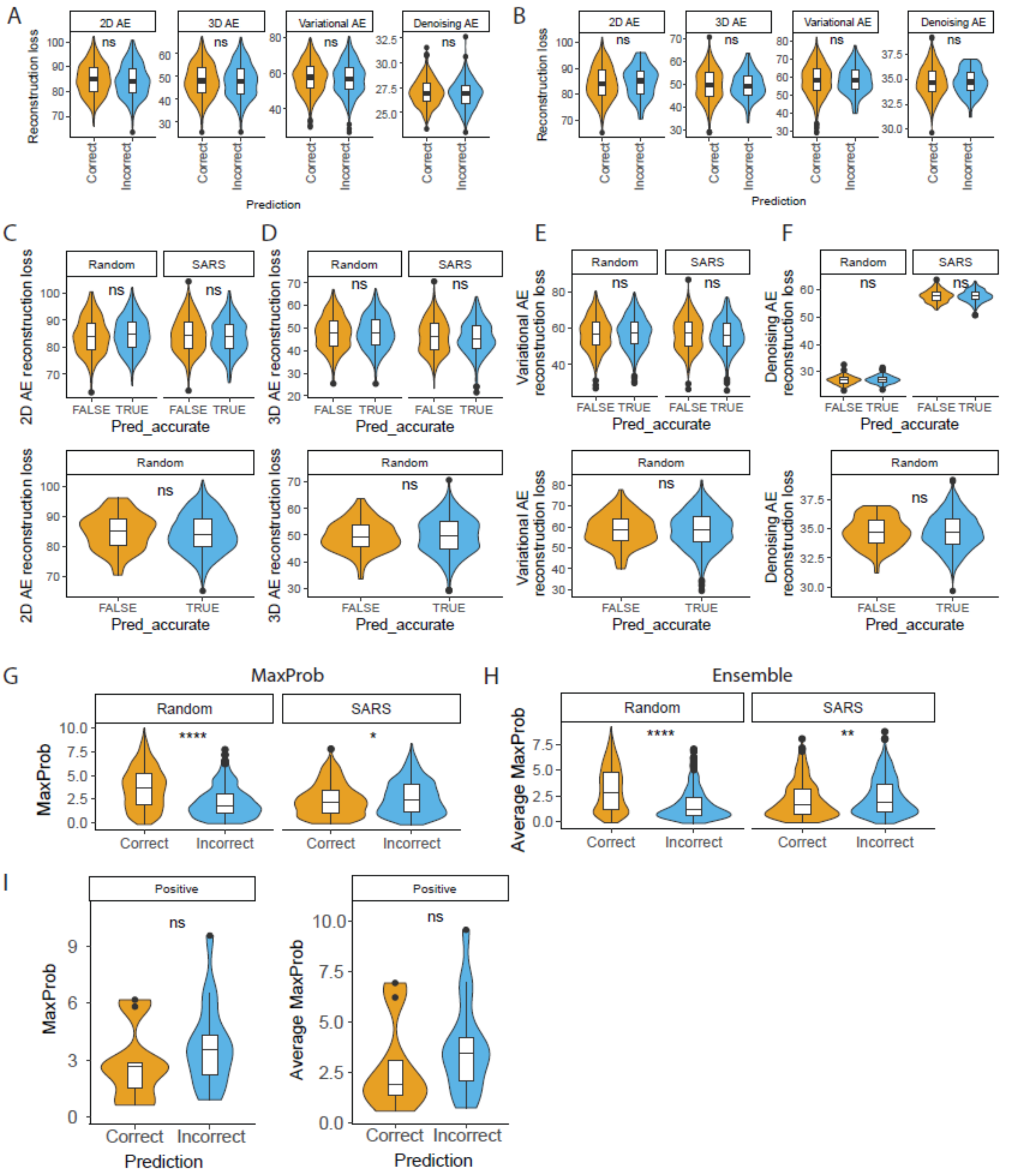


**Supplementary Figure 7. OOD detection.** A-B. Anomaly detection using autoencoders to discriminate correctly or incorrectly predicted pathogenic (A) and self- (B) peptides. Autoencoders included are 2-dimensional autoencoder (2D AE), 3-dimensional autoencoder (3D AE), variational autoencoder and denoising autoencoder. Statistical significance by p-values from Student's t-test. ns: non-significant. We observed no significant difference in reconstruction loss between correctly and incorrectly predicted peptides, implying that the autoencoder-based methods cannot effectively identify OOD inputs. We speculated that unsupervised anomaly detection methods, such as reconstruction loss from autoencoders, were ineffective because ‘species’ was not the primary driver of peptide clustering at contact positions, which was often driven by the cognate TCRs to which they were bound. Likewise, density-based or distance-based methods that use sequence-representation similarity would not be able to detect OODs. C-F. Anomaly detection using autoencoders – 2D-AE (C), 3D AE (D), variational autoencoder (E) and denoising autoencoders (F) for discriminating correctly vs. incorrectly predicted pathogenic (top) and self- (bottom) peptides. For SARS-2 peptides, non-SARS-2 peptides were used to train autoencoders to predict correctly vs. incorrectly predicted SARS-2 peptides. Statistical significance by p-values from Student’s t-test. ns: non-significant. G-H Distribution of maximum softmax probability (MaxProb) (G) and average maximum softmax probability (average MaxProb) from 10 ensembled models (H) for randomly divided data (resemblance of 10-fold cross-validation) and cross-species data (i.e. model trained using non-SARS-CoV-2 peptides to predict SARS-CoV-2 peptides). I. MaxProb (left) and average MaxProb of 10 ensembled models (right) for glioblastoma cancer peptides predicted to be immunogenic using the TRAP self-antigen model.

**Supplementary Tables**

Supplementary Table 1. Pathogenic dataset
Supplementary Table 2. Self-antigen dataset
Supplementary Table 3. Pathogenic TRAP model hyperparameter optimisation
Supplementary Table 4. Self-antigen TRAP model hyperparameter optimisation
Supplementary Table 5. Pathogenic sequence patterns
Supplementary Table 6. Self-antigen sequence patterns
Supplementary Table 7. Pathogenic OOD model calibration metrics
Supplementary Table 8. Self-antigen OOD model calibration metrics
Supplementary Table 9. GBM data
